# Trogocytosis of cancer-associated fibroblasts promotes pancreatic cancer growth and immune suppression via phospholipid scramblase anoctamin 6 (ANO6)

**DOI:** 10.1101/2023.09.15.557802

**Authors:** Charline Ogier, Akino Mercy Charles Solomon, Zhen Lu, Ludmila Recoules, Alena Klochkova, Linara Gabitova-Cornell, Battuya Bayarmagnai, Diana Restifo, Aizhan Surumbayeva, Débora B. Vendramini-Costa, Alexander Y. Deneka, Ralph Francescone, Anna C. Lilly, Alyssa Sipman, Jaye C. Gardiner, Tiffany Luong, Janusz Franco-Barraza, Nina Ibeme, Kathy Q. Cai, Margret B. Einarson, Emmanuelle Nicolas, Andrei Efimov, Emily Megill, Nathaniel W. Snyder, Corinne Bousquet, Jerome Cros, Yunyun Zhou, Erica A. Golemis, Bojana Gligorijevic, Jonathan Soboloff, Serge Y. Fuchs, Edna Cukierman, Igor Astsaturov

**Affiliations:** Program in Cancer Signaling & Microenvironment, Fox Chase Cancer Center, Philadelphia, PA 19111, USA; The Marvin & Concetta Greenberg Pancreatic Cancer Institute, Fox Chase Cancer Center, Philadelphia, PA, 19111, USA; Centre de Recherches en Cancérologie de Toulouse (CRCT), Université de Toulouse, INSERM UMR-1037, CNRS ERL5294, Equipe de Recherche Labellisée “Ligue Contre le Cancer”, Toulouse, France; Fels Cancer Institute for Personalized Medicine, Lewis Katz School of Medicine at Temple University, Philadelphia, PA 19140, USA; Department of Biomedical Sciences, School of Veterinary Medicine, University of Pennsylvania, Philadelphia, PA 19104, USA; Department of Bioengineering, Temple University, Philadelphia, PA, 19140, USA; Molecular and Cell Biology and Genetics (MCBG) Program, Drexel University College of Medicine, Philadelphia, PA 19102, USA; Department of Cardiovascular Sciences, Lewis Katz School of Medicine, Philadelphia, PA 19104, USA; Hôpital Beaujon - INSERM U1149 - Clichy, France; Department of Cancer and Cellular Biology, Lewis Katz School of Medicine, Philadelphia, PA, 19140, USA

## Abstract

In pancreatic ductal adenocarcinoma (PDAC), the fibroblastic stroma constitutes most of the tumor mass and is remarkably devoid of functional blood vessels. This raises an unresolved question of how PDAC cells obtain essential metabolites and water-insoluble lipids. We have found a critical role for cancer-associated fibroblasts (CAFs) in obtaining and transferring lipids from blood-borne particles to PDAC cells via trogocytosis of CAF plasma membranes. We have also determined that CAF-expressed phospholipid scramblase anoctamin 6 (ANO6) is an essential CAF trogocytosis regulator required to promote PDAC cell survival. During trogocytosis, cancer cells and CAFs form synapse-like plasma membranes contacts that induce cytosolic calcium influx in CAFs via Orai channels. This influx activates ANO6 and results in phosphatidylserine exposure on CAF plasma membrane initiating trogocytosis and transfer of membrane lipids, including cholesterol, to PDAC cells. Importantly, ANO6-dependent trogocytosis also supports the immunosuppressive function of pancreatic CAFs towards cytotoxic T cells by promoting transfer of excessive amounts of cholesterol. Further, blockade of ANO6 antagonizes tumor growth via disruption of delivery of exogenous cholesterol to cancer cells and reverses immune suppression suggesting a potential new strategy for PDAC therapy.

## Introduction

Cancer cells are characterized by an increased requirement for lipids to sustain their growth and to promote membrane-based receptor signaling [1, 2]. As *de novo* lipid synthesis consumes energy and oxygen, most aggressive human cancers resort to metabolic parasitism [3], i.e., uptake of cholesterol and other lipids from exogenous sources. While tumors normally acquire lipids and other nutrients from the bloodstream, the PDAC tumor mass is remarkable in its lack of functional intratumoral blood vessels [4], consisting predominantly of dense connective tissue populated by cancer associated fibroblasts (CAFs). How the pancreatic cancer cells obtain their essential nutrients, especially membranous lipids such as cholesterol, has thus been mysterious but of high interest, as determining how PDAC cells acquire exogenous lipids could facilitate a potential therapeutic strategy. To this end, some studies have proposed a direct uptake of lipids from the interstitial fluid, given high levels of larger lipid particles and soluble nutrients can be found in this PDAC compartment [5]. Hence, a continuous stream of exogenous lipids including cholesterol, from blood to the interstitial fluid, is a major impediment for the intended anti-PDAC activity of dietary lipid restriction [6] as well as inhibitors of *de novo* cholesterol biosynthesis such as statins [7, 8]. Yet, the lack of tangible targets and incomplete understanding of this microenvironmental delivery mechanism has been the major obstacle for efficacious restriction of lipids in blood or interstitial fluid experimentally and clinically [4, 5, 9, 10].

This study arose following our prior analysis of PDAC cells unable to produce endogenous cholesterol because of blockade of the endogenous biosynthetic pathway by by genetic knockout of the distal enzyme NSDHL in a PDAC model dependent on mutation of *Kras* and *Trp53* [7]. Strikingly, cell lines derived from *Nsdhl*-deficient tumors rapidly died when cultured in lipid poor media, even though *Nsdhl*-deficient tumors grew very well *in vivo*, implying that these cancer cells were highly capable of extracting the exogenous cholesterol from the *in vivo* milieu [7]. This led us to investigate the mechanism of intratumoral delivery of lipids and the role of CAFs in the process.

In this study, we have discovered that CAFs transfer exogenous cholesterol and other nutrients via a cell contact-dependent transfer of plasma membrane fragments in a process known as *trogocytosis* [11] (trogo = ‘to gnaw’ or ‘to nibble’) whereby PDAC cells ‘nibble’ CAF membranes. We find that trogocytosis is the dominant means by which PDAC cells derive lipids from the tumor microenvironment and have outlined the mechanism by which such trogocytosis is regulated by a phospholipid scramblase anoctamin 6 (encoded by *ANO6* gene). Since ANO6 is highly expressed in CAFs, ANO6 inhibition starves PDAC tumor cells of lipids.

Another feature of the pancreatic cancer microenvironment is the typical high degree of immunosuppression, that has confounded the application of immunotherapies in this cancer type [12-15]. Of note, this immunosuppressive function has also been linked to CAFs, which can inhibit the activity of T cells in pancreatic tumors [12, 14, 16-18]. Extending our studies, we find that CAF-dependent trogocytosis also transfers cholesterol into cytotoxic T cells (CTLs), causing their exhaustion, and inhibition of ANO6 in CAFs also attenuates this CTL immunosuppressive effects. Together, these findings lead us to propose that blockade of ANO6 diminishes the CAF-mediated trogocytic transfer of exogenous cholesterol and may constitute a new effective therapeutic target against PDAC.

## Results

### Pancreatic cancer stroma as a transit station for lipid delivery

To gain insight into the ability of PDAC cells to take up exogenous cholesterol *in vivo*, we determined the time-course and the transit dynamics of blood-borne low-density lipoprotein (LDL) to cancer cells within the pancreatic tumor mass, generated by orthotopic implantation of DsRed-labeled syngeneic murine PDAC cells deficient in *Nsdhl*, an essential cholesterol pathway gene [7] (**Fig. 1a, Supplementary video 1**). Using multiphoton intravital microscopy of these orthotopic pancreatic tumors, we traced the rate of cancer cell uptake of intravenously injected LDL particles containing BODIPY-conjugated cholesterol [19] (green). As the 25 nm-sized LDL particles rapidly enter the blood vessels and permeate to the interstitial fluid, the uptake of LDL by the cells within the pancreatic tumor mass becomes visible only as large intracellular endosomes, containing BODIPY-cholesterol, accumulate [20] (**Fig. 1a, 1b** and **Supplementary video 1**).

**Figure 1.**
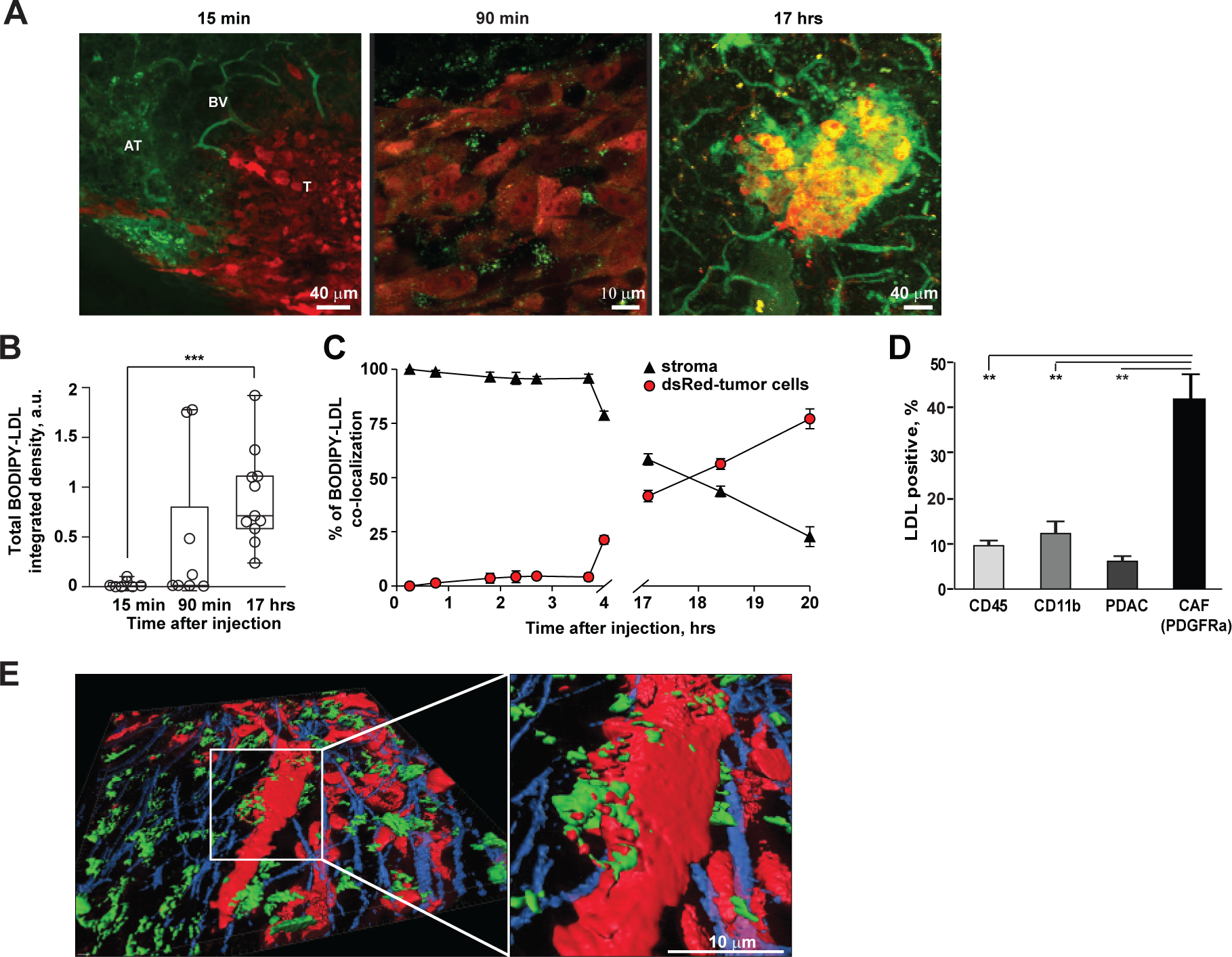
Delivery of low-density lipoprotein particles to pancreatic cancer cells is mediated by cancer-associated fibroblasts. (**A**) Intravital microscopy images of orthotopic DsRed-tagged (*red*) pancreatic tumors at 15 minutes, 90 minutes, and 17 hours after intravenous injection of donor LDL labeled with BODIPY-conjugated cholesterol (*green*). AT, acinar tissue; BV, blood vessel; T, tumor. (**B**) Total integrated intensity of BODIPY-cholesterol over time. Data are represented as mean ±SEM. ***, *p* <0.001 as compared with 15 min post injection (unpaired two-tailed t-test). (**C**) Co-localization of BODIPY-cholesterol with DsRed-expressing carcinoma cells over time. Graph represents pooled data from 2-3 anesthetized animals imaged at 1–2-hour intervals by multiphoton microscopy. (**D**) Flow cytometry enumeration of LDL label in subsets of cells from disintegrated tumors 90 minutes post intravenous administration of fluorescent LDL. Data are represented as mean ±SEM. **, *p*<0.01, ***, *p* <0.001 as compared with CAFs (unpaired two-tailed t-test). (**E**) Enhanced intravital images of endosomal aggregates containing BODIPY-cholesterol and cancer cells in pancreatic tumors. *Red*, KPCN349 murine pancreatic carcinoma cells expressing DsRed; *green*, BODIPY-conjugated cholesterol; *blue*, second harmonic generation collagen. See also Figure S1.

We then performed a more extensive time course, comparing BODIPY signal within DsRed-positive PDAC cells versus unlabeled stroma. Unexpectedly, for up to 4 hours post-injection the exogenously delivered lipids localized exclusively to the stroma, and not to the cancer cells (**Fig. 1c**). We confirmed by flow cytometry analysis at 90 min post-injection that LDL was predominantly detected in PDGFRα-positive cells, the majority of which are known to be cancer-associated fibroblasts (CAFs), but only modest levels were gauged in fluorescently tagged PDAC cells, or in CD11b-or CD45-positive immune cells (**Fig. 1d, S1**). Following this initial delay, BODIPY-cholesterol progressively co-localized to the DsRed-positive pancreatic carcinoma cells, at 17-20 hours post intravenous injection (**Fig. 1c, Supplementary video 2**), whereas the amount of BODIPY-cholesterol in the stroma concurrently decreased to background levels. Most of the BODIPY-positive PDAC cells were in direct physical contact with BODIPY-positive stromal cells, with clearly visible cellular interdigitations (**Fig. 1e**). These results implied that exogenous LDL lipids predominantly transit via CAFs for their ultimate delivery to PDAC cells.

### CAFs provide cholesterol to PDAC cells via a cell contact-dependent mechanism

To determine whether the LDL-cholesterol transfer observed *in vivo* can also be modeled in *in vitro*, we first co-cultured BODIPY-cholesterol labeled pancreatic CAFs with DsRed-positive PDAC cells. This resulted in a rapid (4-12 hours) acquisition of BODIPY signal by PDAC cells (**Fig. 2a**). Importantly, visual enumeration revealed that the ability to acquire BODIPY-cholesterol was largely limited to those PDAC cells that were in direct physical contact with CAFs (**Fig. 2b,c, S2a**). In support of this observation, no transfer was detected between the same cells when these were separated by a Transwell membrane (4 µm pores permeable for soluble molecules and lipid particles; **Fig. 2a** - left panel). Excitingly, detailed microscopic analyses of time-lapse images of PDAC cells revealed synapse-like contacts with CAFs (**Fig. 2c, S2b, Supplementary video 3**), and indicated that PDAC cells “nibble/acquire” plasma membrane (PM) bleb-like protrusions produced by CAFs at the site of the heterotypic cell-cell contacts.

**Figure 2.**
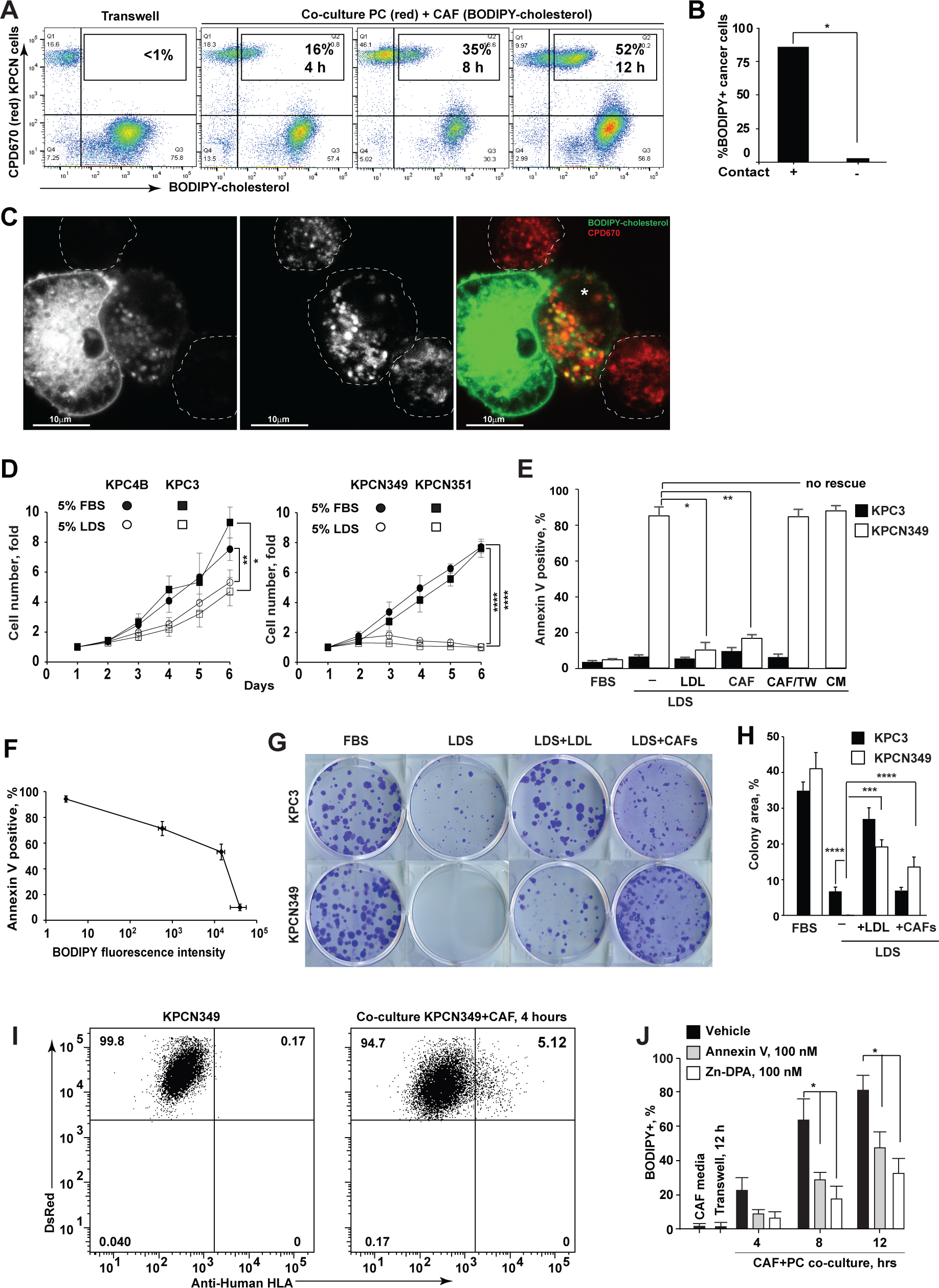
Transfer of membrane cholesterol from cancer-associated fibroblasts to cancer cells via trogocytosis. (**A**) Flow cytometry measurement of uptake of CAFs membranes by KPCN349 cells in co-culture. CAFs were labeled with BODIPY-cholesterol and KPCN349 cells with CPD670 and plated at 1:1 ratio for the indicated times. (**B**) Uptake of BODIPY-cholesterol by KPCN349 cells in co-culture occurs via cell contacts with CAFs (the white-colored asterisk indicates the PDAC cells in direct contact with the CAF cell in this image). Data are represented as mean ±SEM. *, *p*<0.05 as compared with KCN349 cells making no contacts (unpaired two-tailed t-test). (**C**) Representative image of CAF membrane uptake by KPCN349 PDAC cells (*red*, CPD670) and CAFs (*green*, BODIPY-cholesterol). (**D**) Fold change in cell numbers of KPC (*Nsdhl*-proficient) or KPCN (*Nsdhl*-deficient) mouse pancreatic adenocarcinoma cells in the absence of exogenous lipids in 5% lipid-depleted serum (LDS). Data are represented as mean ±SEM. *, *p*<0.05, **, *p*<0.01, ****, *p* <0.0001 as compared with cells grown in 5% FBS (unpaired two-tailed t-test). (**E**) Apoptosis in LDS estimated by Annexin-V/FITC labeling of *Nsdhl*-deficient KPCN349 cells is rescued by co-culture with CAFs, or by addition of 50 µM LDL cholesterol. CAF/TW, co-culture of CAFs and cancer cells in transwell plates; CM, CAF-conditioned 5% LDS/DMEM media for 72 hours. Data are represented as mean ±SEM. *, *p*<0.05,**, *p*<0.01, ****, *p* <0.0001 as compared with KPCN349 cells in LDS (Wilcoxon-test). (**F**) Inverse relationship of Annexin-V-positive carcinoma cells and BODIPY-cholesterol acquired from labeled CAFs in co-cultures. CAFs and KPCN349 cells were plated at varying densities 72 hours prior to Annexin-V labeling. (**G**) 10-day colony formation by mouse pancreatic carcinoma cells in the indicated conditions. (**H**) Quantification of the colony areas as in (C). Data are represented as mean ±SEM. *, *p*<0.05,**, *p*<0.01, ****, *p* <0.0001 as compared with KPCN349 cells in LDS (Mann Whitney-test). (**I**) Flow cytometry measurement of uptake of human HLA by KPCN349 cells in co-culture. Human CAFs were labeled with BODIPY-cholesterol and KPCN349 cells with CPD670 and plated at 1:1 ratio for 8 hours. (**J**) Masking of cell surface phosphatidylserine with recombinant annexin V or bis-Zn-dipicolylamine antagonizes uptake of CAF membranes by KPCN349 cells. CAFs and KPCN349 cells were labeled as in (F). Annexin V and Zn-DPA were added to co-culture media. Data are represented as mean ±SD. *, *p*<0.05 as compared with vehicle (unpaired two-tailed t-test). See also Figure S2.

To further investigate the functional significance of the *in vivo* transfer of cholesterol to PDAC cells, we examined the ability of CAFs to support the viability of cholesterol auxotrophic PDAC cells *in vitro*. In the absence of lipids and cholesterol in the media (growth in lipid depleted serum, or LDS), murine *Kras^mut^Trp53^-/-^* (KPC) cell lines with an intact *Nsdhl* gene had only minor growth delay in LDS medium, and did not undergo apoptosis (**Figs 2d, e**). In contrast, the *Nsdhl*-null KPCN349 and KPCN351 cells did not grow, as anticipated due to the cholesterol auxotrophy imposed by Cre-mediated deletion of this essential cholesterol pathway gene [7] (**Fig. 2d**), and these cells underwent apoptosis after 72 hours (**Fig. 2e**). However, the KPCN PDAC cells were significantly rescued from apoptosis by direct addition of 50 µM cholesterol LDL particles. Importantly, this result was successfully phenocopied by co-culture with human CAFs derived from PDAC patients (**Fig. 2e**).

Cell contact was once again essential for the ability of CAFs to sustain cholesterol-auxotroph PDAC cell survival, as conditioned medium (CM) or co-culture with CAFs separated from PDAC cells by a mesh in Transwell plates, failed to rescue the LDS-cultured *Nsdhl*-null PDAC cells (**Fig. 2e)**. Of note, CAF-generated CM contained measurable concentrations of cholesterol (3-5 µM, **Fig. S2c**). Nonetheless, these levels were insufficient to support the viability of the cholesterol-auxotroph PDAC cells, which required greater than 30 µM LDL cholesterol to maintain PDAC cells viability *in vitro* (**Fig. S2d**). Furthermore, we varied the likelihood of PDAC-CAF cell contacts by altering the CAF-PDAC cell ratio and plating cell density and found an inverse relationship between the uptake of BODIPY-cholesterol in CAF membranes and the percentage of annexin-V positive apoptotic PDAC cells (**Fig. 2f**). Further, to investigate the durability of PDAC dependence on CAFs, we maintained long-term cultures of KPCN349 (cholesterol auxotroph) and KPC3 cancer cells in FBS, LDS, LDS supplemented with 50 µM cholesterol in LDL, and in co-cultures with CAF in LDS-containing media (**Fig. 2g, h**). These clonogenic PDAC cell assays indicated that CAFs provide ample supply of cholesterol to maintain PDAC cell viability and long-term growth.

### PDAC cells obtain cholesterol via trogocytosis of CAF PMs

Trogocytosis is a process by which the cells engage in direct physical contact for the receiver cell to acquire the donor cell’s PM. Trogocytosis has been previously demonstrated for both immune cell synapses [21, 22], and in human development [23]. In CTLs, trogocytosis causes uptake of the human leukocyte antigen (HLA) MHC-I and peptide complexes from antigen-presenting cells [21, 24, 25]. Importantly, upon co-culture of human CAFs with mouse PDAC cells, we observed a similar acquisition and re-expression of human HLA proteins on the murine PDAC surface (**Fig. 2i**).

Exposure of phosphatidylserine (PtdSer) on the outer leaflet of the PM has been established as the “eat me” signal in trogocytosis executed by T cells [25, 26]. Hence, we asked whether blockade of PtdSer antagonizes CAF-to-PDAC cell trogocytosis. Indeed, masking the CAF extracellular exposure of PtdSer, by pre-incubating CAFs with label-free annexin V or zinc-dipicolylamine (Zn-DPA) [27], significantly reduced the uptake of BODIPY-labeled CAF membranes (**Fig. 2j**). These results indicated that PtdSer externalization is important for trogocytosis at heterotypic CAF-PDAC cell contacts.

### ANO6 scramblase confers poor survival in pancreatic cancer

To gain mechanistic insight into regulation of trogocytosis in PDAC, we sought out candidate enzymes responsible for PtdSer externalization in CAFs. Exposure of PtdSer on the outer leaflet of PM is the consequence of lipid redistribution, or ‘‘scrambling”, between the inner and outer leaflets of the PM to be regulated by lipid scramblases [28, 29]. In contrast to Xk-related 8 (XKR8) scramblase, which is involved in the exposure of PtdSer on the external leaflet of the entire PM during apoptosis [30, 31], in non-apoptotic cells phospholipid scramblase anoctamin 6 (ANO6) induces local externalization of PtdSer in response to discrete/localized elevated intracellular calcium under physiologic stimuli [28, 31]. Notably, trogocytosis of CAF membranes by Panc-1 cells was markedly reduced in calcium-free media, or at +4°C (**Fig. S2f**). Taken together, these results suggest that CAF trogocytosis may involve a calcium dependent PtdSer scramblase of the anoctamin family.

Since *ANO*-family paralogs have not been previously implicated in cancer, we examined their expression in mouse [7] and in human [32] pancreatic cancer single-cell RNA sequencing datasets (**Fig. 3**). Of the five *ANO*-family paralogs in mice (*Ano3, 4, 6, 7 and 9*) with experimentally confirmed PtdSer scramblase activity [33], only *Ano6* was expressed in CAFs, endothelial cells and adenocarcinoma cells within the KPC tumor cellular populations as we have previously defined [7] (**Fig. 3a, S3a, b**). Of note, the pattern of ANO6 expression in human PDAC was similar, suggesting a unique and non-redundant function for this scramblase (**Fig. 3b, S3c, d**). Further, high expression of ANO6 protein (**Fig. 3c**) or transcript (**Fig. 3d**) are associated with worse overall survival of patients with stages 1-3 of PDAC as well as in breast and cervical cancers in the TCGA cohorts (**Fig. S3e**). We also assessed the expression of ANO6 by western blot in a panel of fibroblastic cells obtained from surgical pancreatic adenocarcinoma tissues. We detected a consistently higher expression of ANO6 in fibroblastic cells harvested from human PDAC tumor masses (e.g., CAFs) compared to those isolated from adjacent, non-malignant, human pancreatic tissues (**Fig. 3e**). Together, these results support a pro-tumoral role for ANO6 in PDAC and other human cancers, with a potentially important role in pancreatic CAFs.

**Figure 3.**
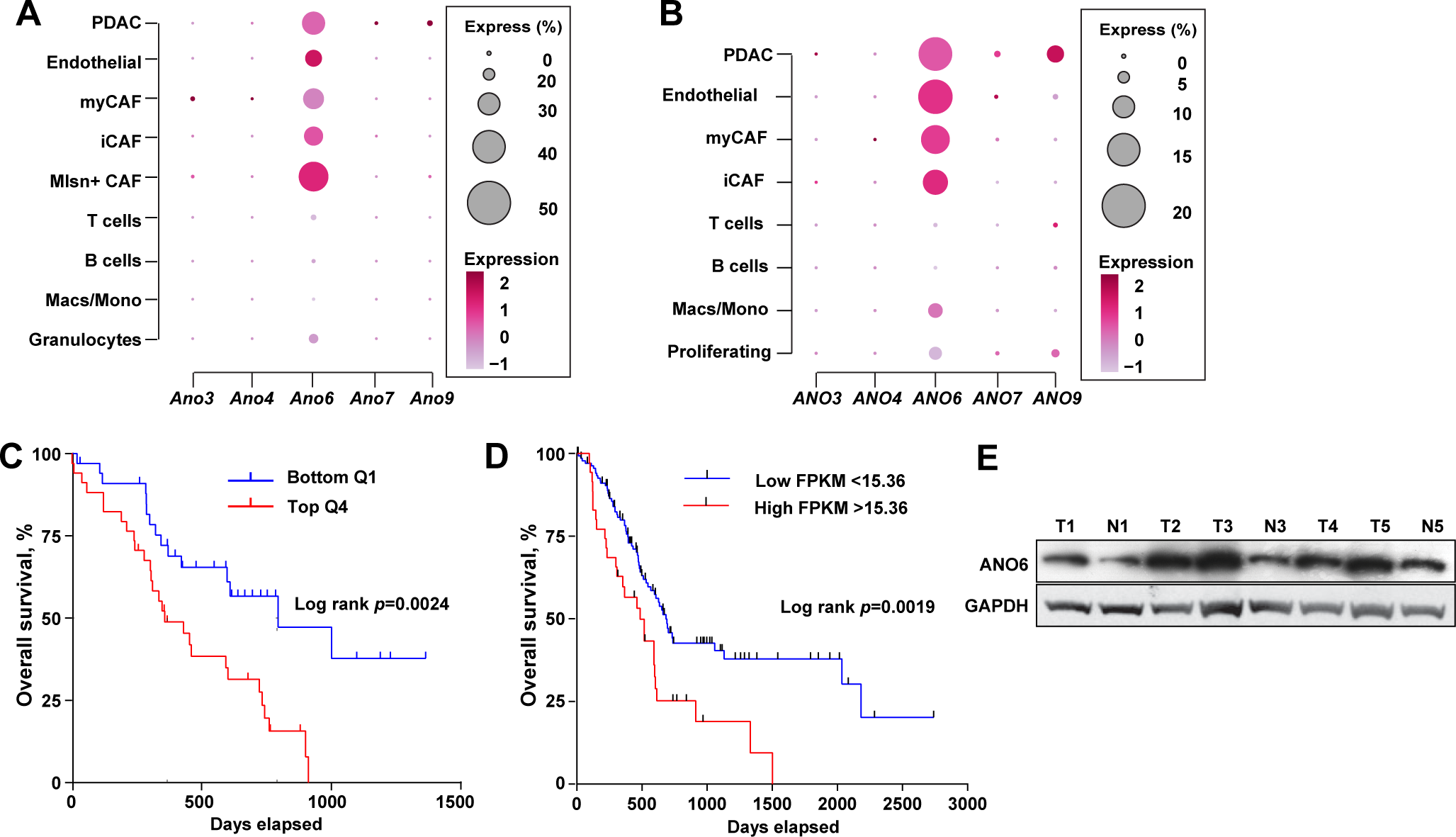
High expression of ANO6 in pancreatic adenocarcinoma confers poor survival. (**A**) *Ano* gene family transcripts expression visualization by single-cell RNA sequencing of 17,205 single cells isolated from four advanced mouse KPC pancreatic tumors. (**B**) *ANO* gene family transcripts expression visualization by single-cell RNA sequencing of 20,929 single cells isolated from five human pancreatic tumors. In A and B, the circle size represents percentage of indicated cells and color intensity represents Z-score normalized expression of the genes. (**C**) Comparison of overall survival of stage 1-3 PDAC patients from the top and the bottom quartiles of ANO6 protein expression obtained from the NCI Clinical Proteomic Tumor Analysis Consortium (CPTAC) dataset. (**D**) Comparison of overall survival of stage 1-3 PDAC patients by *ANO6* mRNA expression above or below the fragments per kilobase of exon per million mapped fragments (FPKM) cut off 15.36. Data was obtained from the NCI TCGA program. In C and D, *p-*values by Mantel-Cox log-rank test. (**E**) ANO6 expression by western blot in protein lysates from fibroblastic cell lines obtained from PDAC tumor (T) or adjacent non-malignant pancreatic tissues (N). See also Figure S3.

### PDAC cells activate ANO6 in CAFs via Orai1-STIM calcium-dependent signaling

To validate the role of ANO6 in CAF membrane trogocytosis, we depleted ANO6 in CAFs using CRISPRi [34] (**Fig. S4a**). As expected, treatment of control CAFs with the calcium ionophore ionomycin increased cytoplasmic calcium and led these cells to externalize PtdSer to the PM. However, ionomycin did not cause externalized PtdSer in ANO6-depleted CAFs (ANO6-KD) or in CAFs pre-treated with the ANO6 inhibitor niclosamide [35] (**Fig. 4a, Supplementary video 4, 5**). Further, co-culture of human CAFs with Panc-1 cells resulted in robust increase in scramblase activity, reflected in a large increase in PtdSer exposed on the CAF PM (**Fig. 4b**). Importantly, we noted that this increase was observed only in control CAFs (ANO6-proficient) engaged in physical contacts with DsRed-tagged Panc-1 cells, but not in ANO6-KD CAFs, CAFs not engaged in physical contacts with PDAC cells, or in CAFs treated with the ANO6 inhibitor niclosamide (**Fig. 4b**). Addition of the PM-impermeable calcium chelator ethylene glycol-tetraacetic acid (EGTA) to control CAFs also prevented ionomycin-induced PtdSer externalization, suggestive of ANO6 inhibition (**Fig. S4b**). Further, quantification of cytosolic Ca^2+^ (as measured by fluorescent calcium sensor Fluo4 (green, **Fig. 4c, d, S4c-e, supplementary video 6**) showed higher levels of cytosolic Ca^2+^ in CAFs in direct contact with PDAC cancer cells (red, **Fig. 4c**), compared to CAFs in monoculture, or to CAFs that did not engage in direct membrane cell-cell contacts with PDAC cells. We concluded that PtdSer externalization by CAFs is largely dependent on calcium-activated ANO6 at heterotypic cancer cell-CAF membrane contacts.

**Figure 4.**
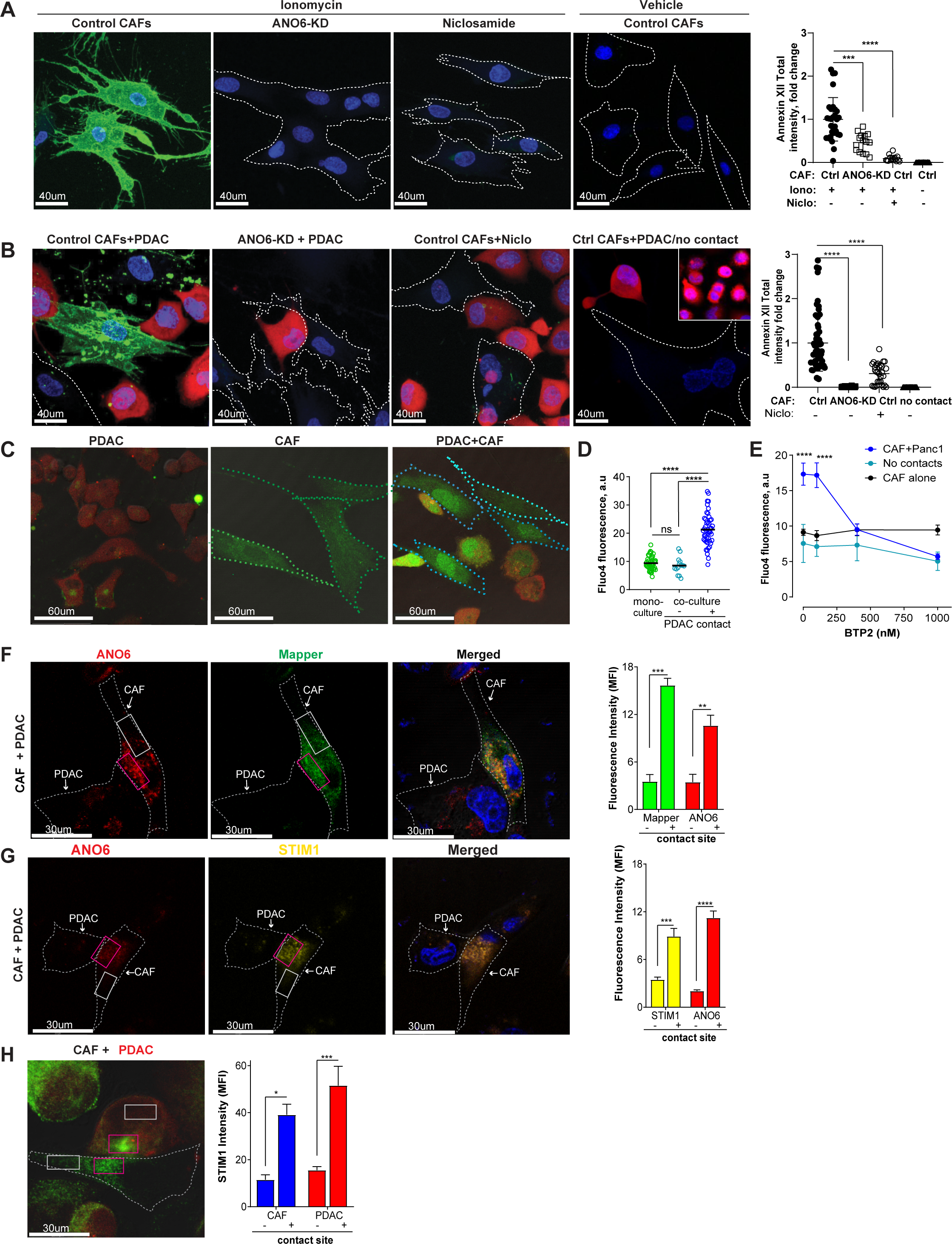
Cancer cells activate ANO6 in CAFs via cellular contact-dependent Orai calcium channels. (**A**) CRISPRi-knockdown of ANO6, or ANO6 inhibitor niclosamide prevents PtdSer externalization induced by calcium ionophore ionomycin (5 minutes at 10 µM) as assessed by intravital annexin XII staining of CAF membranes. *Right*, graph represents fold change of total annexin XII fluorescence intensity relative to control CAFs; *bar*, mean ±SD. *, *p*<0.05, **, *p*<0.01, ****, *p* <0.0001 as compared with control CAFs treated with ionomycin (unpaired two-tailed t-test). (**B**) Co-culture of DsRed-tagged Panc-1 cells (*red*) and unlabeled CAFs induces PtdSer externalization assessed by intravital annexin XII staining of CAF membranes. Note absence of annexin XII labeling in CAFs not making membrane contacts with Panc-1 cells (*outlined*, Ctrl CAF+PDAC/no contact). *Right*, graph represents total annexin XII fluorescence intensity; *bar*, mean ±SD. *, *p*<0.05, **, *p*<0.01, ****, *p* <0.0001 as compared with control CAFs co-cultured with Panc-1 cells (unpaired two-tailed t-test). (**C**) Representative images of fluorescent cytosolic Ca^2+^ indicator Fluo4 in cocultures of DsRed-tagged Panc-1 and CAFs. (**D**) Dot plot represents Fluo4 intensity in individual CAF cells from 3 independent experiments; *bar*, mean ±SEM; *c*olors: *green*, CAFs alone; *cyan*, CAF cells in co-culture not making contacts with Panc1 cells; *blue*, CAFs making membrane contacts with Panc1 cells. *ns*, not significant; **, *p* < 0.01; ****, *p*< 0.0001 as compared with Fluo4 intensity in CAFs making membrane contacts with Panc-1 cells (One-way ANOVA). (**E**) Orai channel inhibitor BTP2 rapidly abrogated CAF-Panc-1 contact-induced Fluo4 signal. CAFs and DsRed-tagged Panc-1 cells were co-cultured as in (D), Fluo4 intensity in CAFs was measured at 5 minutes after addition of indicated concentrations of BTP2. *C*olors: *black*, CAFs alone; *cyan*, CAF cells in co-culture not making contacts with Panc1 cells; *blue*, CAFs making membrane contacts with Panc1 cells. Data are represented as mean ±SD. ****, *p*<0.0001 as compared with CAFs making membrane contacts with Panc-1 cells (unpaired two-sided t-test). (**F**) Reconstituted (maximum intensity) confocal Z-stack images of CAFs transfected with GFP-MAPPER and ANO6-mCherry prior to co-cultured with Panc-1 cells. *Right*, graph represents fluorescence intensity distribution within indicated colored boxes. Data are represented as mean ±SEM. **, *p*<0.01, ***, *p* <0.001 comparing the cell contact box (*red*) with the cell body (*white*) (One-way ANOVA). (**G**) CAFs were transfected with YFP-STIM1 and ANO6-mCherry before co-culture with Panc-1 cell as in (F). Graphs represent STIM1 and ANO6 fluorescent signal intensity distribution within indicated colored boxes. Data are represented as mean ±SEM. ***, *p*<0.001 ****, *p* <0.0001 comparing the cell contact box (*red*) with the cell body (*white*). (**I**) Immunofluorescent labeling of the endogenous STIM1 (green) in CAF and DsRed-tagged Panc-1 cells. Graph represents STIM1 fluorescent signal intensity distribution within indicated colored boxes. Data are represented as mean ±SEM. *, *p*<0.05; ***, *p*<0.001 comparing the cell contact box (*red*) with the cell body (*white*) (One-way ANOVA). See also related Figure S4.

To identify the physiological receptors activating ANO6 in CAFs, following interaction with PDAC cells, we used a pharmacological strategy to probe distinct routes of Ca^2+^ entry into the cytosol. The store-operated calcium entry (SOCE) inhibitors BTP2 [36, 37] and CM4620 [38, 39] (**Fig. 4e, S4c-e, supplementary video 7**) were used to query whether PDAC cells trigger extracellular Ca^2+^ entry into CAFs. This is because this process is known to be mediated by Orai channels (the pore-forming subunit mediating SOCE). Also, phospholipase C inhibitor U73122 was used to determine if contacts with PDAC cells stimulate inositol triphosphate-induced Ca^2+^ release from intracellular stores in the endoplasmic reticulum (ER) of CAFs (**Fig. S4c**). Both BTP2 and CM4620 effectively blocked cytosolic Ca^2+^ elevation in CAFs engaged in heterotypic cell-cell contacts with PDAC cells, and notably these inhibitors also prevented PtdSer externalization in CAFs (**Fig. S4e, S4f**), while U73122 had no effect, suggesting that Orai channels may be activated by PDAC cells in a storage-independent manner.

Based on this findings, we considered the possibility that heterotypic CAF-cancer cell membrane contacts could functionally simulate immunological synapses (IS) [40], in which the Ca^2+^ signaling machinery becomes polarized at sites of cell-cell membrane contacts [41-51]. In an immune synapse, Orai activity is regulated by members of the Stromal Interacting Molecule (STIM) family of ER Ca^2+^ sensors, which physically associate with and activate Orai channels [52]. Typically, STIM1 and STIM2 respond to ER Ca^2+^ depletion by translocating within the ER towards ER-PM junctions. However, during IS formation, STIM translocate towards the IS in a manner not directly dependent upon ER Ca^2+^ concentration [52]. To assess the possibility of a similar occurrence at heterotypic PDAC-CAF interaction sites, the local CAF of ER-PM junctions was marked using the GFP-MAPPER [53], an artificial construct that labels sites of close ER-PM apposition. Based on prior reports, STIM1 should be detected at these ER-PM junctions [54]. Strikingly, CAF GFP-MAPPER puncta were polarized in the direction of heterotypic CAF-PDAC cell-cell junctions in the co-cultures (**Fig. 4f, S4g**), yet these were distributed throughout the entire cell contour in CAF monocultures (**Fig. S4h-k**).

Further, since ANO6 activity is dependent on STIM/Orai, and since STIM is known to modulate the function of numerous PM resident proteins [52, 55, 56], we reasoned these proteins may physically interact at ER-PM contact sites. In support of this concept, transfection of fluorescently labeled STIM1 and ANO6 showed co-enrichment of these proteins at heterotypic PDAC-CAF contact sites (**Fig. 4g**), in contrast to both proteins being evenly distributed in CAF monocultures (**Fig. S4h-k**). A similar localization was observed in cancer cells and CAFs engaged in synapse-like contacts as assessed by a direct labeling of endogenous STIM1 (**Fig. 4h**). Our findings indicate that the functions of STIM, Orai and ANO6 in CAFs are closely coupled, and ultimately, are required for PtdSer externalization to facilitate trogocytosis of CAF PM by recipient cells (e.g., PDAC cells).

### Cholesterol delivery to cancer cells is critical for tumor-promoting function of ANO6 in CAFs

We next determined whether ANO6 expression in CAFs is critical for the ability of these cells to provide cholesterol and thus support the viability of cancer cells in co-cultures under lipid poor media and *in vivo* (as shown above in **Figs. 1** and **2**). Hence, we compared ANO6-KD CAFs to CRISPRi control CAFs modified with non-targeting gRNAs (i.e., ANO6-positive CAFs). ANO6-KD fibroblasts exhibited a slower growth rate (**Fig. 5a**). ANO6 depletion abrogated the ability of CAFs to produce an anisotropic extracellular matrix (e.g., structurally organized ECM) characterized by fibronectin fibers aligned in parallel patterns (**Fig. 5b, c**) [57]. Furthermore, ANO6-KD fibroblasts acquired stellate morphology (**Fig. 5d, S5a**), and accumulated lipid droplets (**Fig. 5d, S5b**), all features of naïve pancreas-resident fibroblastic cells [15, 58]. In contrast to naïve fibroblastic cells (naturally tumor-suppressive [59]), pro-tumoral CAF are activated in a reciprocal manner by their self-generated extracellular matrix and typically express activated matrix-dependent integrin signaling pathway reflected by the phosphorylated focal adhesion kinase (FAK) and AKT [13, 57]. Of note, ANO6-KD CAFs exhibited markedly reduced levels of autophosphorylated focal adhesion kinase (pY397) FAK, and (pS473) AKT (**Fig. 5e**), suggesting that ANO6 inhibition indeed triggers a functional “CAF normalization.”

**Figure 5.**
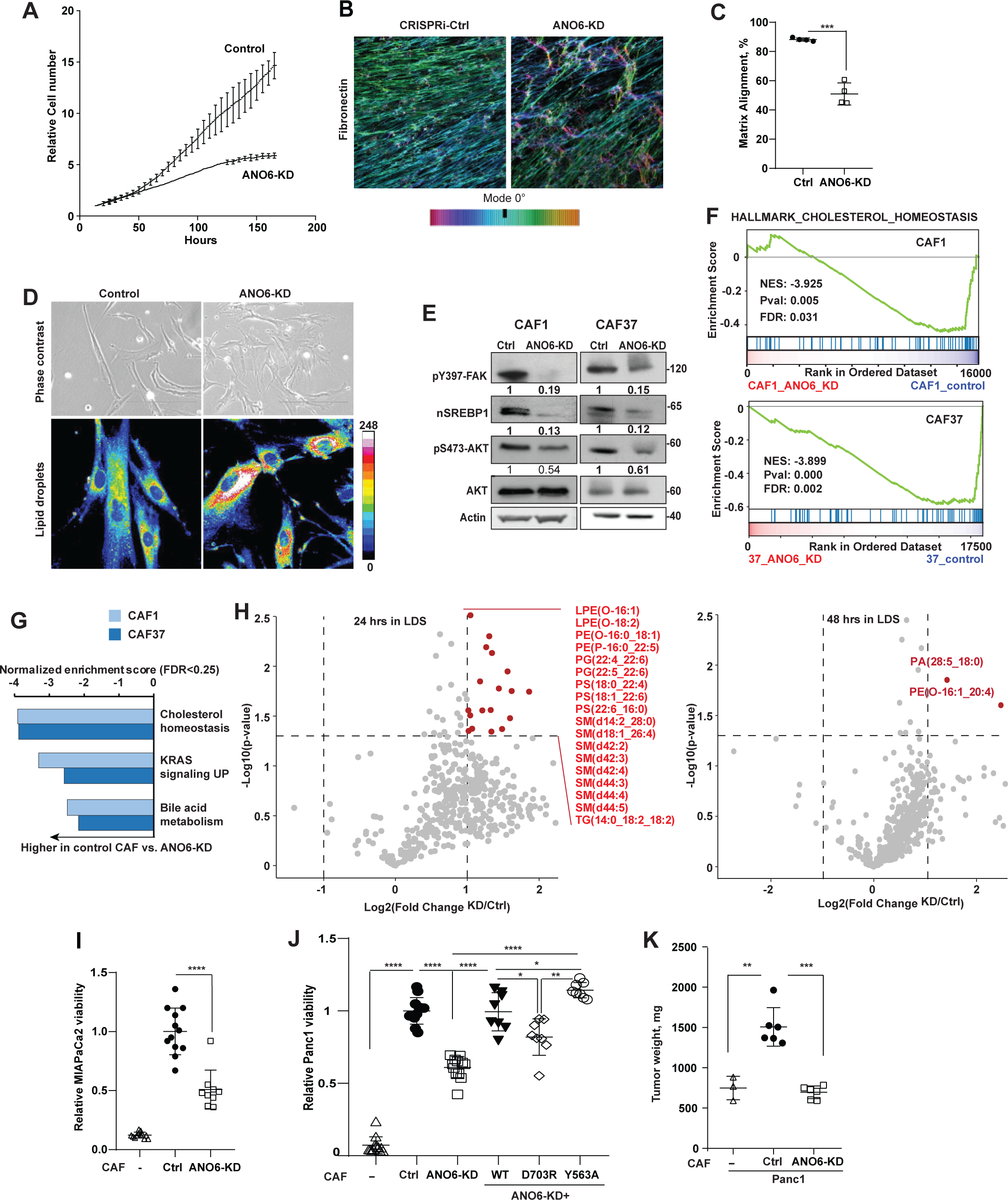
ANO6 regulates cholesterol metabolism and tumor-promoting function of CAFs. (**A**) Growth of ANO6-KD fibroblasts in vitro by continuous xCELLigence impedance assay. (**B**) Representative image demonstrating alignment of extracellular matrix fibers produced *in vitro* by CAF-Control and ANO6-KD fibroblasts. Anti-fibronectin was used to label fibers of extracellular matrix; the fiber angle distribution was determined by ‘Orientation J’ plugin in Image J. Images represent normalized values of fibers angles as shown in the colored bar on the right. (**C**) Percentages of fibronectin fibers aligned at less or equal 15° angles in matrices produced by control CAFs as compared to ANO6-KD CAFs. Graph represents mean ±SD. ***, *p*<0.001 (unpaired two-tailed t-test). (**D**) Representative Images of CRISPRi-control and ANO6-KD fibroblasts in phase contrast (top) or Nile red (bottom) labeling of lipid droplets. (**E**) Expression of indicated protein epitopes in total cellular lysates of CRISPRi-control and ANO6-KD fibroblasts. Bands density relative to control CAF lysates is shown under each lane. (**F**) Normalized enrichment scores by for select (FDR<0.25) Hallmark signatures in ANO6-KD fibroblasts relative to the CRIPSRi-controls. (**G**) Hallmark cholesterol homeostasis signature enrichment plots for ANO6-KD and controls of CAF1 and CAF37 lines of CAFs. (**H**) LC-MS profiling of cellular lipids in ANO6-KD and CRISPRi-control CAFs cultured in LDS for 24 (*left*) and 48 hours (*right*). Volcano plots represent data normalized to cell counts and spiked-in standards in 3 independent replicates of measurement. Lipids with more than 2-fold change (FC) and *p*<0.05 in 3 replicates are marked in red (unpaired two-sided t-test). Lipids abbreviations: LPE, lysophosphatidylethanolamine, PE, phosphatidylethanolamine, PG, phosphatidylglycerol, PS, phosphatidylserine, SM, sphingomyelin, TG, triglyceride. (**I**) Viability of cholesterol auxotroph MiaPaCa^ΔNSDHL^ cells (CRISPRi-depleted of NSDHL) cultured for 4 days in 10% LDS alone, or in the presence of control CAFs, or ANO6-KD CAFs. Data represent fluorescence of DsRed tagged MiaPaCa^ΔNSDHL^ cells relative to co-cultures with control CAFs. Each element represents individual technical replicate. ****, *p*<0.0001 (unpaired two-sided t-test). (**J**) Viability of cholesterol auxotroph Panc-1^ΔNSDHL^ cells (CRISPRi-depleted of NSDHL) cultured for 4 days in 10% LDS alone or in the presence of control CAFs, ANO6-KD, or ANO6-KD modified with lentiviral expression of a full-length murine ANO6-mCherry, or with indicated mutations. Data represent fluorescence of Panc-1^ΔNSDHL^ cells pre-labeled with Hoechst-33342 relative to co-cultures with control CAFs. Each element represents individual technical replicate. *, *p*<0.05, **, *p*<0.001 (unpaired two-sided t-test). (**K**) Tumor weights at 5 weeks following orthotopic implantations of 5x10^5^ Panc-1^ΔNSDHL^ cells alone or with ANO6-modified CAFs at 1:3 ratio. See also Figure S5.

Given the noted lipid droplet accumulation in ANO6-KD CAFs, we next examined the effects of ANO6 on fibroblastic lipid homeostasis. Expression of an exogenous wild type allele of ANO6, but not of the inactive D703R ANO6 mutant (which does not respond to ionomycin [60, 61]) reduced lipid droplets in ANO6-KD CAFs (**Fig. S5c**) suggesting ANO6 is a regulator of lipid turnover in CAFs. The genesis of lipid droplets has been reported as an adaptive mechanism to sequester excessive amounts of lipids to counter lipotoxicity [62, 63]. In the context of ANO6 deficiency, we considered that altered PtdSer polarization in the PM and the intracellular membranes might cause perturbations of trafficking of exogenous lipids from the endocytic routes to the PM and to intracellular organelles [64-66]. Alternatively, lipid droplets could be caused by excessive endogenous biosynthesis of lipids including cholesterol and triglycerides [67]. To distinguish between these two possibilities, we compared transcriptomes and lipid composition of ANO6-KD fibroblasts versus control CAFs (**Fig. 5f-h**). Reduced expression of a nuclear fragment of SREBP1, a transcription factor known for promoting transcription of genes in lipid biosynthetic pathways (**Fig. 5e**), and reduced expression of hallmark cholesterol metabolism signature genes (**Fig. 5f,g**) in ANO6-KD CAFs implied that indeed lipid droplets, in the context of ANO6 loss, are likely the result of an impaired intracellular lipid turnover and not triggered by excessive *de novo* biosynthesis. Further, liquid chromatography mass spectroscopy-based quantification of lipids in ANO6-KD and control CAFs showed increased abundance of multiple phospholipids and triglycerides in the ANO6-deficient fibroblasts (**Fig. 5h**). Preponderance of these lipids in ANO6-KD CAFs (known to be associated with lipid droplets [62]) was significantly reduced after 48 hours of culturing in LDS-containing media, suggesting ANO6 regulates the uptake as well as the processing of exogenous lipids in CAFs. Taken together, these results support the notion that ANO6 regulation of lipids homeostasis and cholesterol metabolism in CAFs may be important for the pro-tumoral function of CAFs via provision of essential exogenous lipids to cancer cells.

In support of this idea, ANO6 inactivation in CAFs significantly reduced the viability of cholesterol auxotroph *NSDHL*-depleted MiaPaCa-2^ΔNSDHL^ (**Fig. 5i**) and Panc-1^ΔNSDHL^ (**Fig. 5j, S5d**) PDAC cells co-cultured with CAFs in LDS-containing media. As anticipated, introduction of the wild-type mCherry-tagged ANO6 to ANO6-KD CAFs not only restored PtdSer externalization in CAFs (**Fig. S5e, S5f**), but also rescued the viability of these cholesterol-auxotroph PDAC cells (**Fig. 5j**). In contrast, co-cultures of Panc-1^ΔNSDHL^ cells with CAFs expressing inactivating D703R mutation in the ANO6 calcium-binding pocket [60, 61] (**Fig. S5e, S5f**) showed lower viability compared to ANO6-KD CAFs that re-expressed wild type ANO6, or a constitutively active conformation of the scramblase, ANO6-Y563A (**Fig. 5j, S5e, S5f**). Importantly, co-implantation of control ANO6^+^ human CAFs with the more aggressive cholesterol auxotroph Panc-1^ΔNSDHL^ human cell model (**Fig. S5g, S5h**) resulted in doubling of orthotopic tumor weights, whereas ANO6-deficient CAFs failed to promote the growth of the xenografts (**Fig. 5k**). These results support the idea that ANO6 plays a critical role in maintaining the growth of cancer cells.

Critically, we also determined that ANO6 blockade antagonized *in vivo* growth of orthotopic grafts of pancreatic cancer cells with intact NSDHL via blockade of lipids delivery. Treatment of orthotopic syngeneic pancreatic tumors derived from the KPC3 cell line with the ANO6 inhibitors niclosamide[35] (**Fig. 6a, b and S6a**) or clofazimine[68] (**Fig. 6a, c, S4f** and **S6b**) significantly reduced tumor weights. To evaluate whether this correlated with cholesterol uptake by cancer cells, we intravenously administered LDL particles carrying BODIPY-cholesterol into niclosamide or vehicle-treated mice bearing DsRed-tagged PDAC KPC3 tumors. Treatment with niclosamide suppressed BODIPY-cholesterol accumulation in the DsRed-tagged PDAC KPC3 cells [7], while the control vehicle-treated tumors showed 6-fold progressive enrichment of BODIPY-cholesterol fluorescence in cancer cells at 16 hours post intravenous injection (**Fig. 6d**). In analyses of intratumoral vesicles containing BODIPY-cholesterol, we also determined progressive cholesterol accumulation in vehicle-treated orthotopic tumors at 19 hours post injection, while a significantly lower accumulation was observed in niclosamide treated mice (**Fig. 6e, f**).

**Figure 6.**
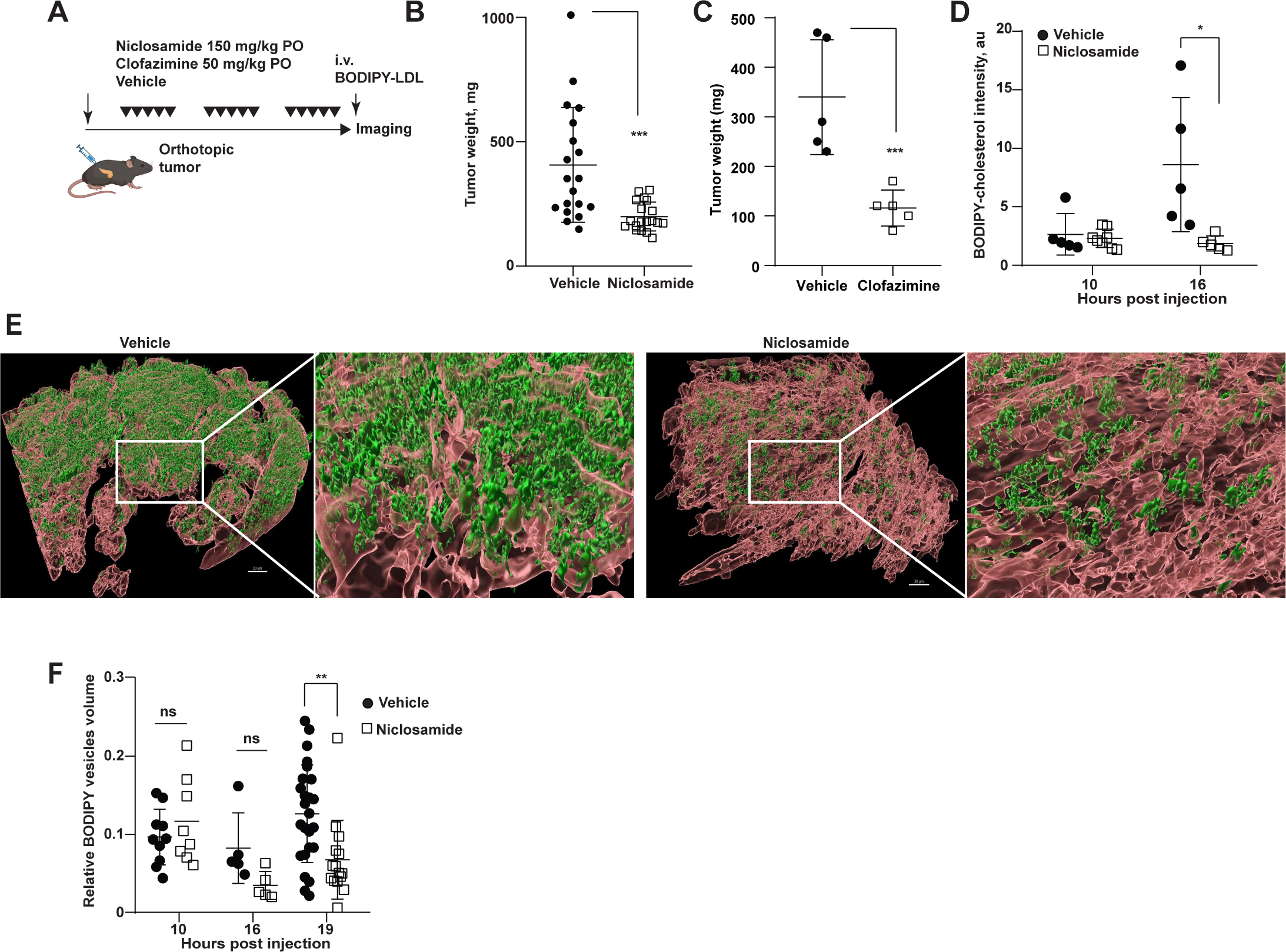
Blockade of ANO6 restrains growth and blocks delivery of exogenous lipids to pancreatic tumors. (**A**) Treatment schema of mice carrying KPC3 syngeneic pancreatic tumor grafts. Animals were given 3 weekly cycles of niclosamide 150 mg/kg/day, clofazimine 50 mg/kg/day, or vehicle followed by intravital imaging and tumor collection. (**B, C**) Weights of orthotopic syngeneic pancreatic tumors following treatment with oral ANO6 inhibitors niclosamide (B) or clofazimine (C) at 3 weeks post implantation. Data are represented as mean ±SD. ***, *p*<0.001 comparing with vehicle-treated mice (unpaired two-tailed t-test). (**D**) Intravital imaging quantification of intratumoral and peri-tumoral BODIPY-cholesterol fluorescence in DsRed-tagged KPC3 tumors at 10, 16 and 19 hours following intravenous administration of labeled LDL. Data are represented as mean±SD of ratios of intratumoral to peri-tumoral BODIPY-cholesterol fluorescence. *, *p*<0.05 (**E**) Representative intravital images of vesicles containing BODIPY-cholesterol within orthotopic DsRed-tagged KPC3 tumors treated with niclosamide or vehicle. *Scale bar*s, 30 µm. (**F**) Quantification of intratumoral vesicles as in (E). Cumulative data from 2 independent imaging experiments are represented as mean ±SD of vesicle volumes enumerated from multiple tumor areas in 2-3 mice at each time point, each element represents vesicles volume in a tumor area. *, *p*<0.05 (unpaired two-tailed t-test). See also Figure S6.

### ANO6-regulated CAF trogocytosis is immunosuppressive against native or CAR-bearing CTLs

Our results so far describe how ANO6 regulates trogocytosis in CAFs. As pro-tumoral CAFs are also known to be highly immunosuppressive (e.g., against CD8-positive CTLs, [12, 14, 16-18]), and because trogocytosis acts as a mechanism by which CTLs acquire the excess of cholesterol and other lipids can trigger lipotoxic ER stress, known to result in CTL exhaustion [21, 24, 69], we investigated if ANO6-regulated CAF membrane trogocytosis also played a role in T cell dysfunction in PDAC.

To this end, we conducted immunoprofiling analyses (**Fig. S7a**) of tumor and splenic tissues harvested from the PDAC tumor-bearing mice treated for 3 weeks with the ANO6 inhibitor clofazimine and compared these to vehicle-treated controls (**Fig. 6c**). We found no clofazimine-induced changes in the frequencies of immune cells in the spleen (**Fig. S7b**). Further, clofazimine also did not change the numbers of intratumoral CD11b^+^F4/80^+^ macrophages, LY6G^+^ granulocytic or LY6C^+^ monocytic myeloid-derived suppressor cells, or the percentage of detected NK cells (**Fig. S7c**). However, PDAC tumors from clofazimine-treated mice included significantly fewer regulatory T cells (Tregs) and increased numbers of intratumoral CD8+ T cells as well as type I conventional dendritic cells (**Fig. 7a**).

**Figure 7.**
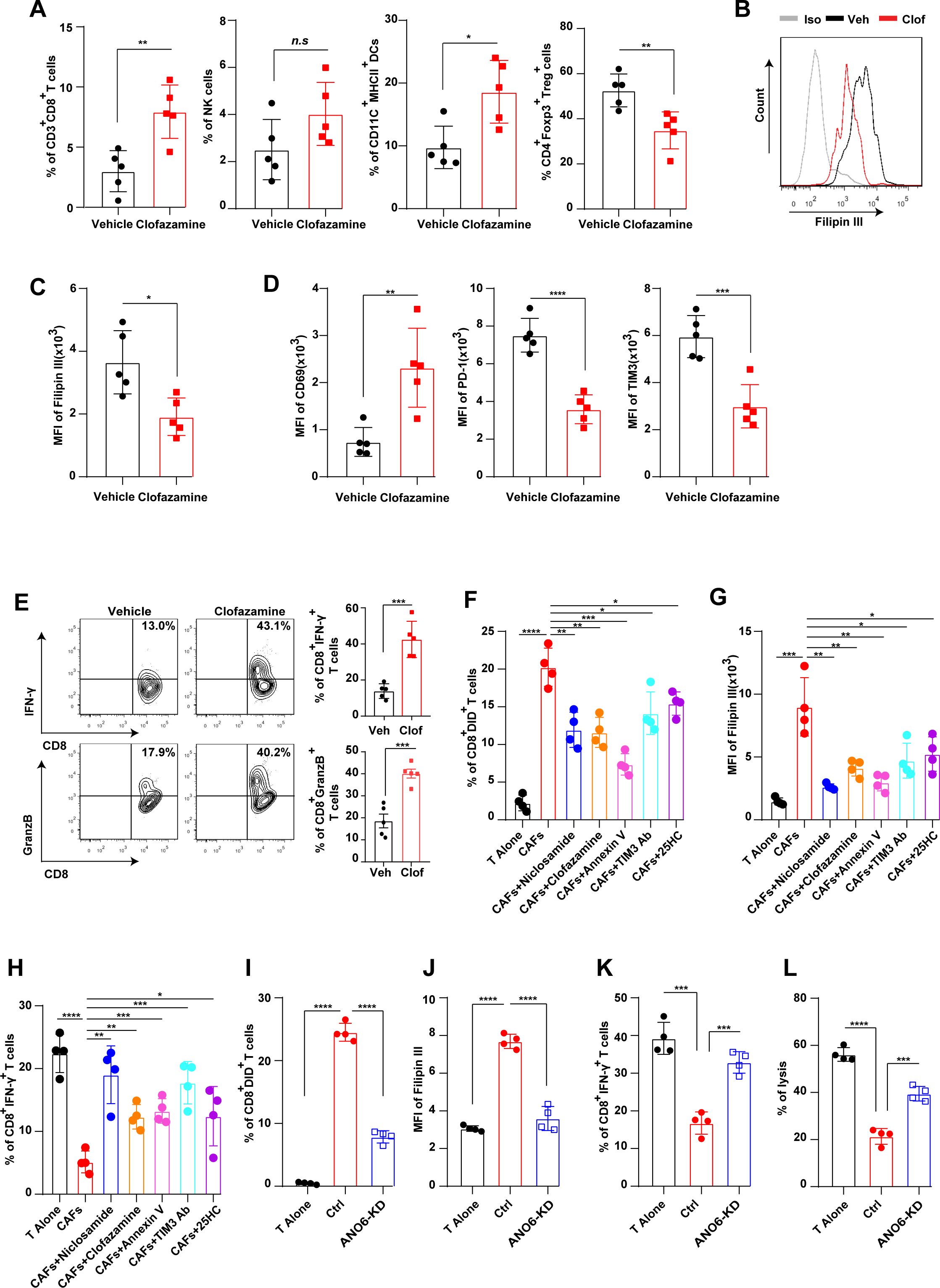
CAF-expressed ANO6 induces the dysfunction of the cytotoxic T lymphocytes in PDAC. (**A**) Flow cytometry percentage of pancreatic intratumoral CD3+CD8+ CTLs, NK cells, conventional type I dendritic cells, and CD4+FoxP3+ regulatory T cells from orthotopic KPC3 pancreatic tumors in mice treated with clofazimine or vehicle as described in Fig. 6C. (**B**) Flow cytometry profiles of Filipin III staining corresponding to cholesterol levels in the intratumoral CTLs from mice treated with clofazimine or vehicle as described in Fig. 6C. (**C**) Quantification of Filipin III mean fluorescence intensity (MFI) related to Fig. 6B. (**D**) Expression of activity (CD69) and exhaustion (PD-1 and TIM3) markers on the intratumoral CD3+CD8+ CTLs isolated from mice treated with clofazimine or vehicle as described in Fig. 6C. (**E**) Effect of clofazimine on intracellular interferon-γ and granzyme B expression in intratumoral CTLs. (**F**) CAFs membrane trogocytosis by OT-1 CTLs. DiD-labeled CAFs were co-cultured with OT-1 CTLs at 1:5 ratio, respectively, in presence or absence of niclosamide, clofazimine, annexin V, anti-TIM3 and 25HC for 12h. (**G**) Enumeration of membrane cholesterol in OT-1 CTLs by Filipin III following co-cultures with CAFs pre-treated as in (F). (**H**) Intracellular interferon-γ in OT-1 CTLs following co-culture with CAFs treatments as in (E). (**I**) CAFs membrane trogocytosis by OT-1 CTLs. DiD-labeled CAF Ctrl or ANO6-KD were co-cultured with OT-1 CTLs at 1:5 ratio for 12 hours. (**J**) Enumeration of membrane cholesterol in OT-1 CTLs by Filipin III following co-cultures with CAF Ctrl or ANO6-KD. (**K**) Intracellular interferon-γ in OT-1 CTLs following co-culture with CAFs. (**L**). Activation of OT-1 CTLs measured by MC38OVA-luc cells lysis following co-culture with Ctrl or ANO6-KD CAFs. In all graphs, data are presented as mean ± SEM from n=5 tumors per treatment or n=4 replicates for each in vitro condition; *p*-values were calculated by unpaired two-tailed t test: *, *p*<0.05; **, *p*<0.01; ***, *p*<0.001; ****, *p*<0.0001. Supplementary figures and data.

We next characterized CD8^+^ CTLs from these mice. Notably and in accord with the hypothesized notion that ANO6 drives pro-tumoral CAF function via regulation of heterotypic cell-cell lipid transfer, clofazimine treatment decreased cholesterol levels in intratumoral CTLs (**Fig. 7b, c**), but failed to do so in splenic CD8^+^ T cells (**Fig. S7d**). This observation suggests that clofazimine can only affect cholesterol levels in T cells that are in direct contact with malignant cells and/or CAFs. Accordingly, clofazimine-induced changes in markers of T cell effector function or exhaustion were not seen in splenic CD8^+^ CTLs (**Fig. S7e**). However, intratumoral CTLs from mice treated with this ANO6 inhibitor displayed increased levels of expression of known T Cell activation markers such as CD69, interferon γ (IFN-γ), and Granzyme B (**Fig. 7d, e**), and decreased expression of known exhaustion markers, such as PD-1 and TIM3 (**Fig. 7d** and **S7f**).

These results suggest that similarly to the heterotypic CAF-PDAC cell contacts, CAF-mediated cholesterol transfer into intratumoral CTLs decreases the anti-tumor function of CTLs. To test this hypothesis and investigate this mechanism in a more defined cellular model, we conducted a trogocytosis assay with CAFs labeled with lipid-soluble DiD dye and co-cultured with OT-1 CTLs carrying a transgenic T cell receptor known to target an ovalbumin epitope. As a control, blockade of trogocytosis with 25-hydroxycholesterol acting on acceptor CTLs [24], effectively inhibited the transfer of DiD from CAFs to CTLs as expected (**Fig. 7f** and **S7g**). Importantly, blockade of PtdSer transfer from CAFs to CTLs, either with a neutralizing antibody against TIM3 (the major receptor for externalized PtdSer and a key regulator of trogocytosis in T cells [25]) or with recombinant Annexin V, significantly reduced CAF PM uptake (labeled by DiD) by CTLs. In support for the critical role of ANO6 in CAF membrane trogocytosis, pre-treatment of ANO6^+^ CAFs with ANO6 inhibitors, clofazimine or niclosamide, notably suppressed the measured transfer of the lipophilic dye (**Fig. 7f** and **S7g**).

Similarly, this blockade of CAF trogocytosis also prevented the increase in the levels of cholesterol in the acceptor CTLs (**Fig. 7g** and **S7h**). These results suggested that CAFs transfer membrane-incorporated cholesterol onto CTLs in an ANO6-PtdSer-TIM3 dependent manner. Importantly, these inhibitors of the ANO6-PtdSer-TIM3 pathways and 25-hydroxycholesterol also reduced the ability of CAFs to inhibit IFN-γ production by the CTLs (**Fig. 7h** and **S7i**). These pharmacological studies were further corroborated in CTL/CAF co-cultures using ANO6-KD CAFs in which reduced DiD uptake (**Fig. 7i** and **S7j**), reduced CTL cholesterol (**Fig. 7j** and **S7k**), increased CTL expression of IFN-γ (**Fig. 7k** and **S7l**), and increased OT-I CTL-dependent lysis of target MC38-OVA cancer cells (**Fig. 7l**), were detected when compared to control, ANO6-expressing CAFs.

Collectively, these results are consistent with our above-postulated hypothesis proposing a key pro-tumoral role for CAF-expressed ANO6, as data suggest that in addition to nurture cholesterol-auxotroph PDAC cells, CAFs drive CAF-to-CTL trogocytosis and thus uphold CTLs immunosuppressed while ensuring high cholesterol levels in both PDAC and CTLs within the pancreatic tumor milieu and resulting in smaller pancreatic tumors (**Fig. 8**).

## Discussion

The unique PDAC microenvironment has long been recognized as nutrient-poor and immunosuppressed, linked to an unusually high proportion of pro-tumoral functioning CAFs to tumor cells (estimated as at least 5 to 1 [70]). In this study, we identify trogocytosis as a versatile mechanism by which CAFs simultaneously support PDAC cancer cell growth and suppress CTL anti-tumor responses. We also show that ANO6 scramblase activity is critical to initiate trogocytosis via exposure of PtdSer on the outer PM leaflet of pro-tumoral CAFs. We report that ANO6 activity is induced by direct heterotypic cell-cell contact (e.g., between PDAC cells and CAFs), which triggers STIM1-Orai signaling at CAF intrinsic ER-PM junctions localized at points of these cell-cell contacts in proximity to ANO6. STIM1, in turn, is known to cause activation of Orai channels [52]. This tripartite association leads to an influx of calcium molecules that is needed to bind and activate ANO6 [28].

Because of the oxygen-poor microenvironment, metabolic parasitism for cholesterol and other nutrients is a common feature of the most lethal basal subtype of pancreatic adenocarcinoma [3, 7], making it a high priority to identify factors essential for this parasitism. For lipids, prior studies have assessed the impact of blocking lipid uptake receptors such as LDLR [71], or scavenge receptors such CD36 [72] and SCARB1 [73]. The efficacy of a single-point blockade in these studies may have been limited by the apparent redundancy of lipid uptake mechanisms, which may underlie the observed cancer cell evasion. An alternative possibility, raised by this study, is that direct trogocytosis-dependent transfer from CAFs to PDAC cancer cells represents the dominant source of uptake for exogenous cholesterol and other lipids carried from blood to the tumor interstitial fluid. To this end, we herein identify ANO6 as a drug amenable and calcium-regulated target with a potential to disable this key mechanism. Our results could also explain the cancer-promoting effect of CAFs in pancreatic desmoplasia, not only in nurturing the cancer cells by providing these with direct access to lipids, but also in repelling the anti-tumor activity of immune cells.

An intriguing element of the mechanistic analysis in this study is that the sustained trogocytic heterotypic cell-cell contacts, between cancer cells and CAFs, are also highly reminiscent of the ones described for immunological synapses [46]. Specifically, these synapses have been described as structured membrane microdomains in which calcium channel, pumps and mitochondria coalesce to generate efficient T cell activation [46]. We were surprised to find that similar synapse-like contacts are formed between the cancer cells and CAFs in which polarized structures including ER-PM contacts sites enriched for STIM and Orai channels mediate contact-dependent calcium influx from the extracellular space to the cytosol. Such contact-dependent calcium signaling via Orai is prerequisite for the scramblase activity of ANO6. Of note, Orai inhibitor zegocractin (CM4260) is being tested in clinical trials of acute pancreatitis (e.g., NCT03401190, NCT04195347; clinicaltrials.gov), a strategy which could be deployed to disrupt the PDAC-CAF heterotypic interactions and CAF trogocytosis.

From this perspective, it is perhaps not surprising that heterotypic CAF-CTL cell-cell interactions exert immunosuppressive effects upon T cells [24], similar to those elicited by membrane transfer from the malignant cells [24]. While CTLs require cholesterol for proliferation and activation [74], we and others [24, 69, 75] have demonstrated that excessive uptake of cholesterol in tumor microenvironment induces exhaustion of CD8-positive T cells. Whereas the role of PtdSer receptor TIM-3 has been previously suggested to play a role in T cell inactivation[25], our new evidence derived from pharmacological and genetic experiments (**Fig. 7, S7**) indicate that ANO6 and externalized PtdSer directly regulate the immunosuppressive function of CAFs [14], which is the focus of this study. Since ANO6 is also expressed in cancer cells, it is likely that a similar trogocytosis-mediated mechanism could also antagonize CTLs that are directly engaged in heterotypic cell-cell contacts with PDAC cells [21, 24].

Finally, antimicrobials such as niclosamide and clofazimine, initially discovered as blockers of COVID19-induced syncytia [35, 68], were effective, among other known targets, in photocopying ANO6 loss in CAFs and in abrogating PtdSer externalization, including controlling the growth of PDAC xenografts *in vivo*. Due to their fibroblastic normalizing effect, evidence by inhibition of CAF-mediated delivery of lipids to both nurture cancer cells and uphold immunosuppression, we propose that these ANO6 inhibitors, albeit indirect yet also well tolerated and inexpensive may constitute future highly valued anti-cancer therapies, especially for CAF-relevant solid tumors such as pancreatic cancer.

## Materials and methods

### Animal Studies

All animal protocols were reviewed and approved by the Institutional Animal Care and Use Committee of Fox Chase Cancer Center. C57BL/6J mice were purchased from Jackson Laboratory and maintained at the FCCC Animal Facility. C.B-17.icr SCID mice were purchased from Taconic, NY.

For the xenograft study, anesthetized 6-8-week-old C.B-17.icr SCID mice (Taconic, NY) were injected to the tail of pancreas with 5x10^5^ of NSDHL-depleted Panc-1 cells (CRISPRi-mediated knockdown) in 30 µL of serum-free DMEM. For co-implantation experiments, 5x10^5^ Panc-1 cells were mixed with 1.5x10^6^ of CAFs in 30 µL of serum-free DMEM.

Syngeneic 5*10^5^ KPC3 cells were orthotopically injected to the tail of pancreas of anesthetized C57Bl/6J mice. Treatment with ANO6 inhibitors niclosamide or clofazimine commenced on 2 days post-implantation: niclosamide (150 mg/kg of mouse body weight, Cayman #10649), clofazimine (50 mg/kg, MedChem Express), or corn oil (50 mg/kg, Sigma#C8267) as vehicle were given by oral gavages 5 days/week for 3 weeks. Mice were kept under defined-flora pathogen-free conditions at the AAALAC-approved Animal Facility of the Fox Chase Cancer Center, Philadelphia, PA. Mice of both genders, equally distributed, were used for the experiments.

### In vivo fluorescent low density lipoprotein imaging by multiphoton microscopy

Mice with established orthotopic tumors roughly 21 days post implantation were anesthetized with 1.5-2% isoflurane/oxygen mixture maintained for the duration of imaging. Pancreatic tumors were exposed through a 1 cm incision on the left flank of the mouse placed on a 35°C heating pad (Kent Scientific). Intensity thresholds were set up to distinguish signal from background fluorescence, and areas positive for DsRed, indicative of tumor cells, were selected for 3D confocal imaging using a 25x objective in the Leica SP8 Dive multiphoton imaging system (Leica Microsystems), prior to the intravenous administration of 150 µL of LDL carrying BODIPY-cholesterol and at 0.5, 1, 5, 16, and 19-hour interval post injection with XY resolution 0.867 um/px and Z distance 2 µm. For each time interval, 2-3 mice were imaged for 0.5-2 hours under continuous anesthesia. At the end of each imaging session, the animals were euthanized, and tissues were collected. The acquired 3D stacks were analyzed using Imaris 10.0.0 software (Oxford Instruments, UK).

For measurements of BODIPY-cholesterol uptake by tumors in Figure 6D, tumor volumes were defined using “surface option” in Imaris package with surface grain size 0.867 µm, intensity threshold and size exclusion 300 voxel to exclude non-specific speckles. Volume outside of the tumor within 20 µm of its boundary was defined using “distance transform” option in Imaris software. Cholesterol volume was identified using “surface option” in Imaris with surface grain size 0.867 um, automatic intensity threshold and size exclusion set at 10 voxel. Mean intensity of cholesterol volume co-localized with tumor volume or volume outside of tumor was measured and ratio inside intensity/outside intensity was calculated.

For measurements of BODIPY-cholesterol vesicles in tumors in Figure 6E, tumor volumes were defined as above. The volumes of BODIPY-cholesterol-positive vesicles were defined using a surface grain size of 0.1 µm and background subtraction with 3 µm sphere, automatic intensity threshold and exclusion of objects below 26 voxels. The volume of vesicles inside the tumor were expressed as a percentage relative to tumor volume.

### Cell Lines

PDAC cells were obtained from ATCC and maintained in DMEM supplemented with 10% v/v FBS, 2mM L-glutamine and 100mg/ml Penicillin/Streptomycin. Murine pancreatic carcinoma cell lines were isolated as reported [7]. All cell lines were regularly tested for Mycoplasma, as determined by PCR detection methods, and all lines tested negative. Human CAF1 cell line was derived from the residuals of PDAC surgical samples in accord with the Fox Chase Cancer Center IRB-approved informed consent from patients who donated samples for research purposes as previously described [76, 77]. An additional similarly harvested human pancreatic CAF line (referred to as CAF37) was obtained from Dr. Corinne Bousquet, Centre de Recherches en Cancérologie de Toulouse, Toulouse, France.

### Functional assessment of fibroblastic/ECM functional units; fibroblast-derived 3D matrix cultures

Functional fibroblastic cell 3D culture units are needed to ensure the pathophysiological signal reciprocity between fibroblastic cells and self-generated ECM is sustained *ex vivo.* These units were generated using our published methods [57, 76, 77]. Briefly, confluent cultures of human pancreatic fibroblastic cells, harvested from surgical samples and immortalized via hTERT overexpression, were supplemented daily with freshly-prepared ascorbic acid (50 ug/mL) for a period lasting 5 days. The resulting fibroblastic/ECM 3D functional units were used for the phenotypic characterization of the units and to report on the pro-tumor vs. tumor-suppressive (normalized) statuses of the experimental fibroblastic/ECM units. For this, lysates obtained from the assorted units were resolved using SDS-PAGE and traits were gauged via western blot; testing for constitutive integrin and PI3K activities depicted by phosphorylated FAK and AKT respectively.

Of note, the quality of ex vivo 3D functional units was measured via ECM thinness (indicative of intact fibrillogenesis), while loss of ECM anisotropy (e.g., low parallel ECM fiber alignment) indicated effective unit normalization; as published [77]. Briley, the latter was measured via indirect immunofluorescence of fibronectin fibers analyzed using monochromatic 2D image projections. Images were uploaded to ImageJ (SCR_003070) and ECM fibers were analyzed with the OrientationJ plugin SCR_014796 http://bigwww.epfl.ch/demo/orientation/. Numerical fiber orientation/angle outputs were normalized by setting each image’s mode angle to 0°, thus correcting angle spreads/distributions to fluctuate between -90 and 90. Angle distributions, for each experimental condition, corresponding to a minimum of three experimental repetitions and five image acquisitions per condition, were plotted and their standard deviations calculated. The percentage of fibers oriented at 15 degrees from the mode (ranging from -15° to 15°) was determined for each image-obtained data to inform the presented graphs.

### Plasmid transfections and CRISPRi-mediated gene silencing

Transfection of GFP-MAPPER (117721, Addgene, Watertown, MA), mCherry-ANO6 (62554, Addgene, Watertown, MA) or STIM1-YFP (19754, Addgene, Watertown, MA) were done by mixing 2.5 µg of plasmid DNA with 7.5 µl of TransIT-X2 reagent (Mirus, Madison, WI). The mixture was added to 10^5^ CAF or PDAC cells cultured in serum-free and antibiotic-free media overnight.

For CRISPRi, we used an all-in-one lentiviral CRISPRi plasmid containing a nuclease-dead Cas9 (dCas9) fused to the transcriptional repressor domain KRAB and a gene specific gRNA[34]. The top 3 best predicted gRNA were chosen from the published study[78] and individually cloned into the all-in-one lentiviral vector, CRISPRi-Puro modified from the Addgene plasmid #71236 (a gift from Charles Gersbach[79] containing a ‘‘stuffer’’ at the gRNA cloning site). Lentiviruses were generated by transfecting the CRISPRi-Puro plasmid, along with the packaging plasmids psPAX2 (a gift from Didier Trono; Addgene plasmid #12260) and pMD2.G (VSV-G envelope, a gift from Didier Trono; Addgene plasmid #12259) with X-tremeGene9 transfection reagent (#6365787001, Sigma-Aldrich) into 293T cells in serum free/antibiotic free media overnight. The following day, media was replaced with complete media containing FBS and media containing lentiviruses were collected at days 2 and 4 post-transfection. The lentiviral media was then filtered through a 0.45 uM filter (#HAWP14250, Millipore) and used to transduce target cells, by culturing cells in lentiviral media plus 10 ug/mL Polybrene (#sc-134220, Santa Cruz Biotechnology). After 24 hours, lentiviral media was replaced with complete media and cells were selected 48 hours later with puromycin (specific for each cell lines) for 14 days. Surviving cells were subsequently expanded and knockdown of target genes was confirmed at the protein and transcriptomic levels.

### Generation of ANO6-mCherry mutant CAFs

Mutations in the ANO6 sequence of plasmid ANO6-Plvx-mCherry-c1 (Addgene #62554) were created using the Q5® Site-Directed Mutagenesis Kit (NEB E0554S) based on the established functional scramblase activity data [60, 61]. Primers were designed with the NEBaseChanger tool (https://nebasechangerv1.neb.com/). The intended substitution for each mutation was confirmed by Sanger sequencing (Azenta) and the mutated plasmids were transfected into HEK293T cells. The resulting viruses were filtered and transduced into CAF ANO6 CRISPRi-depleted cells. After growth and expansion, cell lines were selected by flow cytometry for mCherry expression.

### Analyses of protein expression by western blotting

Cultured cells were lysed in RIPA buffer (#24928, Santa Cruz) with phosphatase inhibitor (#1862495, Thermo Fisher Scientific) and protease inhibitors (#1861278, Thermo Fisher Scientific) on ice and cleared by centrifugation (15 min at 17,000 g). The protein concentration was measured with a Pierce BCA Protein Assay Kit (#23225, Thermo Fisher Scientific). Proteins were separated on 4-12% Bis-Tris Protein gels (Invitrogen) and then horizontally transferred to the Immobilon-FL PVDF membrane (#IPFL00010, Millipore). Primary and secondary antibodies were used at the concentrations indicated below according to manufacturer’s instructions. The density of obtained bands was quantified with Image J software (FIJI).

### Lipid droplet fluorescence measurements; normalized CAF assessment

Quantification of lipids droplets accumulated in PDAC or CAFs cells (5000 cells per well in a 96-well plate) was conducted using the Lipid Droplets Fluorescence Assay Kit (Cayman #No. 500001 or AAT Bioquest #22730) according to the manufacturer’s instructions. The fluorescence intensity, indicative of lipid droplet accumulation, was measured using a plate reader, also following the manufacturer’s recommended settings.

### Cholesterol measurement

Cholesterol level in PDAC or CAFs cells were measured by Amplex Red Kit (#A12216, Life Technologies). Briefly, 500,000 cells were seeded in a 6-well plate and cholesterol was measured in cell lysate or supernatant according to the manufacturer’s instructions. BCA test was performed in parallel for normalization.

### Phosphatidylserine externalization assay

CAFs were seeded in 24-well glass bottom plates (#81156, Ibidi, Fitchburg, WI) at a density of 20,000 cells per well. Cells were then treated with niclosamide (1 μM) for 2h. After one wash with FBS-free media at RT, cells were incubated with 250 μl of 1:100 annexin XII and 1:200 propidium iodide (pSIVA Abcam #ab129817), with or without 10 μM ionomycin. Cells were then incubated (5 min 37C) and imaged by fluorescence microscopy using a Nikon A1 camera (Nikon) linked to a Nikon Eclipse Ti2-E Inverted Microscope Imaging System (Nikon) for each channel. Fifteen images per well were acquired.

### Colony formation assay

10^3^ PDAC cells were plated in 2.5 ml of media on 6-well plates with or without 10^5^ CAFs in DMEM supplemented with 10% v/v FBS and 2mM L-glutamine with 100µg/ml Penicillin/Streptomycin. After 24 hours, the old media was replaced with new media according to experimental conditions. After 10 days, the cultures were washed with PBS twice and fixed in 10% methanol 10%/acetic acid solution for 15 minutes at RT and stained for 20 minutes with 0.4% Crystal Violet in 20% ethanol. Plates with colonies were scanned and images were analyzed with ImageJ software using the Colony Area plugin (Guzman et al., 2014).

### Annexin V apoptosis and viability assays

PDAC cells were pre-labeled with eFluor670 (CPD670) according to the manufacturer’s instructions (#65-0840-85, eBioscience) and plated either alone or in co-cultures with CAFs at 10^5^ cells in 2 ml per well in 6-well plates in DMEM supplemented with 10% FBS, 2 mM L-glutamine, and 100 µg/ml Penicillin/Streptomycin. After 24 hours, the medium was replaced with indicated experimental media supplemented with 10% FBS, 10% lipid-depleted serum (LDS), or LDS supplemented with purified donor LDL (#360-10, LEE Biosolutions, Maryland Heights, MO, USA). After 72 hours, cells were harvested by trypsinization and stained with Annexin V-FITC (#640906, Biolegend) or Annexin V-PE (#640908, Biolegend) according to the manufacturer’s instructions. Dead cells were determined by DAPI staining (#D9542, Sigma-Aldrich), fluorescence was measured using BD LSRII flow cytometer and analyzed with FlowJo software. For direct enumeration of viable cells in co-cultures with CAFs, we seeded CAFs at 12,000 cells per well in 96-well plate (#3603, Corning) in DMEM/10% FBS. The next day, the media was replaced with 10%/LDS/DMEM and 4,000 DsRed-tagged Panc-1 cells were plated on the top of CAFs at 3:1 ratio. After 96 hours, fluorescence of viable DsRed-positive Panc-1 cells was measured using Spark Multimode Microplate Reader (Tecan) with parameters set at 540 nm for excitation and 590 nm for detection. To measure the dynamics of cell proliferation in vitro, we used xCELLigence E-Plate (ACEA BioSciences,) as per the manufacturer’s recommended protocol. Impedance values were recorded every hour. Data was graphed using GraphPad Prism software.

### Trogocytosis assays in vitro by flow cytometry

To label cellular membranes, pairs of lipophilic fluorescent dyes were used according to the manufacturer’s instructions in the following combinations: 1) adherent CAFs were stained with 1 µM PKH67 (Sigma-Aldrich), Panc-1 were stained in suspension with 10 µM CellTracker™ orange CMTMR (Thermo Fisher Scientific); 2) CAFs labeled with 0.5 µM BODIPY-cholesterol (Topfluor Cholesterol, #DO-016545, Avanti Polar Lipids) dissolved in 0.1 mM methyl-beta-cyclodextrin (#C4555, Sigma) with 2 µM cholesterol (#C8667, Sigma-Aldrich) in MEM media at 37°C for 30 min in suspension; murine or human (Panc-1) PDAC cells were stained in suspension with CPD670 (#65-0840-85, Thermo Fisher Scientific). After labeling, CAFs were co-incubated for the indicated time intervals with PDAC cells in 12-well culture plates at ratio of 2:1 in 0.5X DMEM media. Co-cultures of CAFs and PDAC cells for 3 minutes on ice minutes were used as negative controls. In experiments assessing the effect of extracellular calcium on trogocytosis, cells were co-cultured in Phosphate Buffered Saline (PBS) containing calcium and magnesium (D8662, Sigma-Aldrich) or in PBS without calcium and magnesium (D8537, Sigma-Aldrich). After co-cultures, cells were collected and pelleted by centrifugation (5 min, 1500 rpm), resuspended in 10% formalin (Sigma) for 10 min at RT, washed and re-suspended in PBS/1% BSA. Trogocytosis was measured as the acquisition of CAF-specific fluorescence in PDAC cells by flow cytometry on MACSQuant® VYB Flow Cytometer (Miltenyi Biotec) or FACScan analyzer (BD Biosciences). Analysis of dual staining was done using FlowJo software (10.8.1, BD Biosciences). Live cells were gated based on FSC/SSC parameters, then single cells were gated based on SSC-A/SSC-H.

For interference with trogocytosis, CAFs were pre-treated prior to co-cultures for 2h with recombinant label-free 100nM Annexin-V (#ab157342, Abcam) or 100 nM Zn-DPA (Molecular Targeting Technologies) and subsequently maintained in co-cultures at the same concentrations.

For trogocytosis between OT-1 and CAF cells, T cells were isolated from splenocytes of OT1 mice (strain 021880, The Jackson Laboratory) and stimulated with OVA_257-264_ (1 μg/ml) for 24 hours as described [24]. CTLs were labeled with DID (DiIC18(5); 1,1′-dioctadecyl-3,3,3′,3′-tetramethylindodicarbocyanine, 4-chlorobenzenesulfonate salt, D7757, Invitrogen) and then co-cultured with CAFs at 5:1 ratio in the presence of 0.05% DMSO as diluent or indicated agents: niclosamide (1 μM), clofazimine (1 μM), anti-TIM3 Ab (4 μg/ml, clone RMT-3-23, BioXCell, Lebanon, NH), or 25-hydroxycholesterol (4 μM, #H1015, Sigma-Aldrich) for 12 hours followed by flow cytometry analysis.

### Tumor dissociation and immune subset analyses

Orthotopically implanted syngeneic KPC3 tumor tissues were harvested and digested with 1 mg/mL Collagenase D (Roche, Switzerland) plus with 100 μg/mL DNase I (Roche) in RPMI medium for 45 min with continuous agitation at 37°C. The digestion mixture was passed through a 100 μm cell strainer to prepare single cell suspension and washed with PBS supplemented with 2mM EDTA and 1% FBS. Single cells were stained with antibodies purchased from BioLegend (San Diego, CA) diluted at 1:200 for staining: CD45-BV785 (cat#103149), CD3-APC (cat#100236), CD8-APC/Cy7 (cat#100714), TIM3-BV421 (cat#134019), CD69-FITC (cat#104506), PD-1-BV605 (#135220), CD11C-PE/Cy7 (cat#117318), MHCII-APC (cat#107615), CD11b-Percp/Cy5.5 (cat#101228), F4/80-BV421 (#123132), Ly6C-BV605 (cat#128036), Ly6G-APC/Cy7 (cat#127624), NK1.1-FITC (cat#156508), CD3-APC/Cy7(cat#100222), CD8-PE/Cy7 (cat#100722), IFN-γ-APC (cat#505810), Granzyme B-Percp/Cy5.5 (cat#372212), CD3-BV421 (cat#100228), CD4-PE/Cy7 (cat#116016), CD25-APC/Cy7(cat#, 101908), Foxp3-AF647 in permeabilized cells (cat#,320014).

### Analysis of cholesterol level and cytokines in T cells

Cholesterol abundance in cell membranes was determined with Cholesterol Detection Assay Kit (Cayman, cat# 10009779). Dissociated cells were fixed with fixative solution for 10 minutes, washed and labeled with Filipin III as per the manufacturer’s instructions. Labeling with Filipin III was done by incubating in the dark for 30 minutes. After two washes for 5 minutes each, labeling was analyzed by flow cytometry (excitation:340-380, emission: 385-470). The IFN-γ, Granzyme B and Foxp3 were detected using Foxp3/Transcription Factor Staining Buffer (00-5523-00, Thermo Fisher Scientific). Cells were stained for cell surface markers at 4°C for 30 min followed by fixation and permeabilization buffer for 30 mins and two washes with permeabilization buffer and incubation with the indicated antibodies for 30 mins, washed with PBS twice followed by flow cytometry analysis.

### Measurement of cytosolic calcium using Fluo4 fluorescence microscopy

CAF (5x10^3^) and PDAC (10^4^) cells were incubated (separately or in co-culture) on coverslips for 12 hrs before loading with the calcium indicator Fluo4-AM (5 µM; Invitrogen, Eugene, OR) in DMEM supplemented with HEPES and 0.5 % FBS and incubated at 37°C, 5% CO_2_ for 30 min. Cells were washed in DMEM before incubating for an additional 30 minutes at 37°C, 5% CO_2_ to permit de-esterification of the dye. Fluorescence emission at 506 nm in response to excitation at 494 nm was monitored for 60 cycles at a frequency of 0.33 Hz using a Leica SP8 laser scanning microscope (Buffalo Grove, IL). Intracellular Ca^2+^ measurements are shown as absolute fluorescence obtained from 35-45 cells obtained using Leica LASX software. Averaged results± SEM from at least three independent experiments were analyzed by one-way ANOVA with GraphPad Prism. BTP2 (STIM/Orai inhibitor; Tocris Bio-Techne Corporation, Minneapolis, MN), CM4620 (STIM/Orai inhibitor; MedChemExpress, Monmouth Junction, NJ) and U73122 (PLC inhibitor; AdipoGen, San Diego, CA) were added to the cells at various concentrations for 10 min to determine their effect on cytosolic Ca^2+^ levels.

### SOCE Measurement in CAFs

CAF (5x10^3^) cells were incubated on coverslips for 12 hrs before loading with the calcium indicator Fura2-AM (2 µM; Invitrogen, Eugene, OR) in cation-safe solution (107 mM NaCl, 7.2 mM KCl, 1.2 mM MgCl_2_, 11.5 mM glucose, 20 mM Hepes-NaOH, 1 mM CaCl_2_, pH 7.2) for 30 min at 24°C as previously described [80]. Cells were washed, and dye was allowed to de-esterify for a minimum of 30 min at 24°C. Cells were treated with Vehicle control, 5µM CM462, or 1µM BTP2 along with 2 μM Thapsigargin for 10 minutes. After Ca^2+^ levels returned to their baseline, 1 mM Ca^2+^ was added. Ca^2+^ measurements were made using a Leica DMI 6000B fluorescence microscope controlled by Slidebook Software (Intelligent Imaging Innovations; Denver, CO). Fluorescence emission at 505 nm was monitored while alternating between 340 and 380 nm excitation wavelengths at a frequency of 0.67 Hz; intracellular Ca^2+^ measurements are shown as 340/380 nm ratios obtained from groups (20-25) of single cells.

### STIM1 Immunocytochemistry

CAF and PDAC cells were co-cultured onto 12-mm circular coverslips before fixing with paraformaldehyde (3.7%; 10 min). Cellular permeabilization was conducted using Tween20 (0.5%; PBS). Samples were blocked with bovine serum albumin (0.5%) before labeling with anti-STIM1 monoclonal antibody (1:50 dilution; 30 min; Sigma-Aldrich, Rockford, IL). Cells were then washed before the addition of Alexa Fluor 488-conjugated goat anti-mouse IgG (1 µg/ml dilution; 30 min; Invitrogen, Eugene, OR). Fixed and labeled cells were mounted onto slides before visualization with a Leica sp8 confocal laser scanning microscope and analysis with LASX software. Data were analyzed by two-way ANOVAs.

### Timelapse image acquisition and analysis

Images were acquired using an ImageXpress micro confocal microscope (Molecular Devices, San Jose,CA) equipped with environmental control, at 10X magnification (10X, Plan Apo) in widefield mode. Four sites per well with 100 uM between images in both the X and Y direction were acquired, in four wavelengths (transmitted light, CY5 (DRAQ5), FITC (SYBR Green), and Texas Red (DsRed). Total timelapse duration was 96 hours, images acquired at 3-hour intervals for a total of 33 timepoints. Images were subsequently analyzed using the ‘mutiwavelength scoring’ module (MetaXpress) for image segmentation and quantitation. Image segmentation gated for positive cells using both size and intensity cutoffs. Scoring profiles for single positive (either FITC positive/DsRed negative or FITC negative/DsRed positive) or double positive cells (FITC positive/DsRed positive) were reported for nuclei positive cells (Total cells = nuclei count or W1). Image acquisition was controlled via MetaXpress (64 bit) version 6.7.0.211 (Molecular Devices). Data was exported to MS-Excel for further analysis.

### In vivo cholesterol transfer measurement by flow cytometry

Tumor dissociation was performed using a gentleMACS Tumor Dissociation Kit (Miltenyi Biotec, Order No. 130-096-730) according to the manufacturer’s instructions. Briefly, each tumor was isolated from the animal in a sterile environment and washed in PBS. A piece was taken for histopathology analysis. The rest of the tumor tissue was placed in the dissociation enzyme mix and minced quickly to get pieces 2mm^3^ in size. Then the enzyme-tissue mixture was transferred into the gentleMACS C tube and incubated at 37C with constant rotation for 40 minutes. After that the tissue was further mechanically processed by the gentleMACS Dissociator. A single-cell suspension was obtained by passing the tissue mixture through a 70mm cell strainer. Dead cells were subsequently removed by the Dead Cell Removal Kit (Miltenyi Biotec, Order No. 130-090-101). Cell lineage was determined with the lineage markers: CD45 (1:200, #103107, BioLegend), CD11b (1:200, #101235, BioLegend), FAP (1:50, #ab28244, Abcam), EPCAM (1:200, #118212, BioLegend) and PDGFRa (1:80, #135905, BioLegend) in the presence of FcR antibody (1:50, #101301, BioLegend) prior to using other antibodies. Live cells were selected based on propidium iodide staining (#421301, BioLegend). Fluorescence detection and sorting were performed with a BD FACS Aria II flow cytometer.

### Analysis of cellular lipids using LC-MS

Optima grade methanol, water, acetonitrile, methyl tert-butyl ether, and 2-propanol were from Thermo Fisher Scientific (Pittsburg, PA). Gasses were supplied by Airgas (Philadelphia, PA). Glassware and HPLC vials were from Waters Corp (Milford, MA).

*Lipid extraction*. Frozen cell pellets (∼1[×[10^6^ cells) were collected in low retention Eppendorf tube and mixed with 0.6 mL 80% methanol (MeOH) and 20 µL of lipidomics internal standard mix (1:1, SPLASH® LIPIDOMIX #330707: Ceramide/Sphingoid Internal Standard Mixture I #LM6002, both from Avanti Polar Lipids, Alablaster, AL) and kept in dry ice. Samples were pulse sonicated for 30x half-second pulse on ice and kept on ice for 20 min for metabolites extraction. Each tube was then vortexed 3x 30 seconds each. The cell homogenates were moved to a 10 mL glass Pyrex tube with screw cap. The Eppendorf tubes were rinsed with 0.5 mL methanol and added to the same glass tube. 5 mL methyl tert-butyl ether (MTBE) was added to each of the tubes and then tubes were shaken vigorously for 30 min. 1.2 mL water was added to each tube and vortexed for 30 sec each. Centrifugation for 10 min at 1000xg created two phases. The top clear phase was moved to a clean glass Pyrex tube and dried down under nitrogen. 200 µL MTBE/MeOH (1:3 v/v) per 10 mg tissue was used to re-suspend the residue. The sample was spun down at 10,000 x g for 10 min at 4°C and only 100 µL were transferred to a HPLC vial for LC-MS analysis. A pooled sample was created by mixing 20 µL of each re-suspended sample and ran as quality control (QC) every 15 samples. 2 µL injections were made in both positive mode and separately in the negative mode.

### Liquid chromatography high resolution -mass spectrometry (LC-HRMS) for lipids

Lipid analysis on the extracted samples was conducted by liquid chromatography-high resolution mass spectrometry as previously described[81]. Briefly, separations were conducted on an Ultimate 3000 (Thermo Fisher Scientific) using an Ascentis Express C18, 2.1 × 150 mm 2.7μm column (Sigma-Aldrich, St. Louis, MO). The flow-rate was 0.4 m /min, solvent A was water:acetonitrile (4:6 v/v) with 0.1% formic acid and 10 mM ammonium formate and solvent B was acetonitrile:isopropanol (1:9 v/v) with 0.1% formic acid and 10 mM ammonium formate. The gradient was as follows: 10 % B at 0 min, 10 % B at 1 min, 40 % B at 4 min, 75 % B at 12 min, 99 % B at 21 min, 99 % B at 24 min, 10 % B at 24.5 min, 10 % at 30 min. Separations were performed at 55 °C.

For the HRMS analysis, a QE Exactive-HF mass spectrometer (Thermo Fisher Scientific) calibrated within 3 days was used in positive ion mode with a HESI source. The operating conditions were: spray voltage at 3.5 kV; capillary temperature at 285°C; auxiliary temperature 370°C; tube lens 45. Nitrogen was used as the sheath gas at 45 units, the auxiliary gas at 10 units and sweep gas was 2 units. Same MS conditions were used in negative ionization mode, but with a spray voltage at 3.2 kV. Control extraction blanks were made in the same way using just the solvents instead of the tissue homogenate. The control blanks were used for the exclusion list with a threshold feature intensity set at 1e10^^5^. Untargeted analysis and targeted peak integration was conducted using LipidsSearch 4.2 (Thermo Fisher Scientific) as described[82]. All samples were analyzed in a randomized order in full scan MS that alternated with MS2 of top 20, with HCD scans at 30, 45 or 60 eV. Full scan resolution was set to 120,000 in the scan range between *m/z* 250–1800. The pool sample was run every 15 samples. Lipids quantification was done from the full scan data. The areas were normalized to the internal standard (SplashLipidoMix Mass Spec Standard, Avanti Polar Lipids, Alabaster, AL) and cell number.

Data analysis was done in MetaboAnalyst 5.0, where the raw peak intensities associated with each metabolite were analyzed using one factor statistical analysis. The processed data were organized into groups that were compared and graphed onto volcano plots to show significantly upregulated and downregulated lipid species.

### Analyses of ANO6 mRNA and protein expression in human cancers

Data for mRNA expression and patient outcomes for human pancreatic ductal adenocarcinoma, cervical cancer and breast cancer were extracted from the publicly available NCI TCGA program. ANO6 protein expression data were obtained from the NCI Clinical Proteomic Tumor Analysis Consortium (CPTAC) dataset. Analyses to compare the overall survival among the ANO6-high and ANO6-low were performed by comparing the Kaplan-Meyer survival curves with log-rank tests. These were calculated using the R ‘survival’ package [83].

### Single-Cell RNA sequencing using 10X Genomics platform

For single-cell RNA sequencing, KPPC mice with advanced tumors (n=3), were sacrificed at 7-8 weeks of age. Single-cell suspensions were isolated from minced tumors using Miltenyi Biotec gentleMacs dissociator in gentleMACS C Tubes (#130-093-23) and mouse tumor tissue dissociation kit (#130-096-730) as per the manufacturer’s instructions. Dead cells were removed by use of Dead Cell removal microbeads (#130-090-101), hematopoietic cells were depleted by CD45 MicroBeads (#130-052-30). The Chromium controller was used to make single-cell droplet with GEM bead. Single-cell suspensions were converted to barcoded scRNA-seq libraries by using the Chromium Single Cell 30 Library, Gel Bead & Multiplex Kit and Chip Kit V3 (10X Genomics, #PN-1000092), loading an estimated 6-12*10^3^ cells per library per the manufacturer’s instructions. Samples were processed using kits pertaining to the V3 barcoding chemistry of 10x Genomics. For each replicate, all tumor samples were processed in parallel in the same thermal cycler. The final libraries were profiled using the Bioanalyzer High Sensitivity DNA Kit (Agilent Technologies) and quantified using the Qubit 2.0 (Thermal Fisher, #Q32851). Each single-cell RNA-seq library was sequenced twice in two lanes of HiSeq 4000 (Illumina) to obtain single-end, 98 bp, approximately 500 million reads per library.

### Bioinformatics of mouse single-cell RNA sequencing

For analyses of human PDAC single-cell RNA sequencing, publicly available Peng et al. (2019)[32] data for samples CRR034503, CRR034504, CRR034505, CRR034506, and CRR034507 were obtained from Genome Sequence Archive under project PRJCA001063. Single-cell RNA-seq data for advanced KPC mouse pancreatic adenocarcinoma were generated by this study. Sequencing data for each sample were processed using CellRanger 7.1.0 (10x Genomics) to align and quantify sequencing reads using a mouse reference genome (GRCm38). Individual count tables were merged using CellRanger *aggr* function. Subsequent data analysis was carried out in R 3.5.1 and the Seurat package (v 3.0.2) as described[7]. We applied the following filters to exclude beads without cells and dead cells by imposing a threshold of at least 500 transcripts measured per cell, and less than 5% mitochondrial reads were set as a threshold to exclude dead cells. We used 20 principal components for dimensionality reduction via UMAP with default parameters. Clusters of cells were identified based on a shared-nearest neighbor graph between cells and the smart moving (SLM) algorithm (resolution = 0.1). Markers for each cluster were identified by reducing the number of candidate genes to those genes which were (i) at least log (0.25)-fold higher expressed in the cluster under consideration compared to all other clusters and (ii) expressed in at least 10% of cells in the cluster under consideration. For genes passing those criteria, significance between cells in the cluster versus all other cells was calculated using model-based analysis of single-cell transcriptomics (MAST) [84] and adjusted with the Benjamini–Hochberg method. Differentially expressed genes were used as input for gene set expression analysis.

### RNA sequencing of CAFs

Human patient-derived immortalized CAFs (lines #7 and #37) modified to express either a non-targeting guide RNA (Control) or gRNA targeting ANO6 promoter (ANO6-KD) and dCas9-KRAB constructs (CRISPRi) grown at 100% confluency in 6-well plates were used. Total RNA was isolated using RNeasy Mini Kit (#74104, Qiagen). mRNA libraries were formed by poly-A capture at Novogene and sequenced in a 150 bp paired-end mode using Illumina NovaSeq platform. Reads were aligned to human genome (hg38 version) using TopHat2 [85] and absolute gene counts were quantified using HTSeq [86]. The resulting gene counts were used as input for differential expression analysis between the control and ANO6-KD CAF cells using DESeq2[87]. Genes that are differentially expressed were selected for subsequent downstream analysis for identification of biological pathways and ontologies using Gene Set Enrichment Analysis (GSEA) (p-value < 0.001)[88] using MsigDB datasets[89]. To graphically represent the significantly enriched datasets (False Discovery Rate < 25%) Enrichment scores and plots were used[90].

### Quantification and statistical analysis

For analysis of continuous data, we used Wilcoxon tests, Mann-Whitney and Student t-test as indicated, and binary outcomes were compared using Fisher’s exact test. Repeated measures (i.e. multiple measures within a single mouse) were analyzed using generalized linear regression models with Generalized Estimating Equations (GEE) [66]. Growth curves were modeled using linear regression with interactions between treatment and time, again using GEE to account for within-sample correlation. Survival time outcomes were assessed using Kaplan-Meier curves with log-rank tests. The statistical details of experiments can be found in the figure legends, figures and text of the Results.

## Supplementary figures and data

**Supplementary Figure 1 related to Figure 1**. (**A**) Gating strategy to enumerate LDL uptake in orthotopic pancreatic tumors. Approximately 90 minutes post intravenous injection via retroorbital vein of 150 µL of fluorescent LDL particles, animals were euthanized and single-cell suspension obtained using Miltenyi Biotec GentleMacs dissociator in gentleMACS C Tubes (#130-093-23) and mouse tumor tissue dissociation kit (#130-096-730) as per the manufacturer’s instructions followed by washes and filtering of non-cellular debris. The cells were first gated using FSC-A and SSC-A plot to eliminate small non-cellular particles and aggregates. LDL-positive population (∼10%) was selected based on a cut off for vehicle-injected negative controls. GFP-or DsRed-positive carcinoma cells were separated from CD45+ and CD11b+ (myeloid/immune) and PDGFRa+ (CAFs) populations. (**B**) Summary table of 3 independent experiments enumerating the percentage of cell types positive for LDL signal.

**Supplementary Figure 2 related to Figure 2. Transfer of membrane cholesterol from cancer-associated fibroblasts to cancer cells via trogocytosis. (A)** Quantification of PDAC-CAF cell contacts duration in co-cultures. Data are compiled from two independent time-lapse microscopy experiments. Contacts durations are inferred from the number of sequential 15-minute frames during which the contacts were maintained. The contact duration data were grouped at 2-hour intervals for simplicity. (**B**) 3D reconstruction of a Z-stack images of CAF (green) and KPCN349 (pink) interface taken at 2-second intervals. *Arrow*, emerging CAF membrane protrusion towards the body of carcinoma cells (trogocytosis). (**C**) Quantification of cholesterol concentration (µM) in 10% fetal bovine serum (FBS) in DMEM and in 10% LDS/DMEM media conditioned with CAFs for 3 days. Data are represented as mean ±SEM of independent experiments. (**D**) Viability of cholesterol auxotroph cell line KPCN349 (NSDHL-null) in 10% LDS supplemented with varying concentrations of cholesterol added to 72 hours culture in the form of human donor LDL. (**E**) The effect of cell plating density on the rate of BODIPY-cholesterol uptake by KPCN349 cells in 24 hours. (**F**) The effects of temperature (*left panel*), or extracellular calcium and magnesium (*right panel*) on CAF membrane trogocytosis by Panc-1 cells co-cultured with CAFs in 10% FBS DMEM (*middle panel*), or in CAF-conditioned media (CM, *middle* panel). Adherent CAFs were stained 1 µM PKH67 (Sigma-Aldrich), Panc-1 were stained in suspension with 10 µM CellTracker™ orange CMTMR (Thermo Fisher Scientific) and added to co-cultures with CAFs for 3 hours. The uptake of PKH67 by Panc-1 cells was analyzed by flow cytometry.

**Supplementary Figure 3 related to Figure 3. Transcriptome analyses of ANO6 expression in mouse PDAC models and human cancers. (A)** Cell type determination based on uniform manifold approximation and projection (UMAP) embedding of transcriptomes of 17,205 single cells isolated from 4 KPPC advanced tumors using the 10X Genomics single-cell RNA sequencing platform. Nine cell types were identified by graph-based clustering are indicated by color. (**B**) Heat map of differentially expressed genes for each cluster. Z-score normalized expression of the enriched genes for each cluster is shown as a log_2_-fold change in cells within a cluster relative to all other cells in the dataset. (**C**) UMAP-embedding of transcriptomes of 20,929 single cells from 5 human pancreatic adenocarcinoma tumors from Peng J. et al. (PMID: 31273297). Raw sequencing data for CRR034503-CRR034507 were downloaded from https://ngdc.cncb.ac.cn/gsa/, accession project CRA001160, and processed as described [1]. Nine cell types were identified by graph-based clustering are indicated by color. (**D**) Heat map of differentially expressed genes for each cluster. Z-score normalized expression of the enriched genes for each cluster are shown as a log_2_-fold change in cells within a cluster relative to all other cells in the dataset. Abbreviations: iCAF, inflammatory cancer associated fibroblasts; myCAF, myofibroblasts; Msln+CAF, mesothelin-positive CAFs. (**E**) The effect of ANO6 transcript expression on overall survival of all human cancers (*left*, TCGA Pan-Cancer bulk RNA sequencing dataset), or selectively in breast invasive carcinoma (*middle*, TCGA BRCA bulk RNA sequencing dataset), and cervical carcinoma (*right*, TCGA CESC bulk RNA sequencing dataset).

**Supplementary Figure 4 related to Figure 4. Influx of extracellular calcium via Orai channels activates ANO6 in CAFs. (A)** Western blot demonstrating loss of ANO6 in CRISPRi-modified CAF1 cells. (**B**) Total annexin XII fluorescence intensity measured by microscopy of viable CAFs pulsed for 5 minutes with 1 µM ionomycin in DMEM containing Ca2+/M2+ or with added cell-impermeable 2 mM EGTA. *Right*, representative images of annexin XII labeling. Data of individual images obtained in two independent experiments are represented as mean ±SD. ****, *p*<0.0001 (unpaired two-tailed t-test). (**C**) *Left panel*, Orai channel inhibitor CM4620 rapidly abrogated CAF-Panc-1 contact-induced Fluo4 signal. CAFs and DsRed-tagged Panc-1 cells were co-cultured as in Fig. 4D, Fluo4 intensity in CAFs was measured at 5 minutes after addition of indicated concentrations of CM4620.*Right panel,* phospholipase C-type phosphatidylinositol-specific inhibitor U-73122 does not abrogate CAF-Panc-1 contact-induced Fluo4 signal. CAFs and DsRed-tagged Panc-1 cells were co-cultured as in Figure 4D, Fluo4 intensity in CAFs was measured at 5 minutes after addition of indicated concentrations of U-73122.*C*olors: *black*, CAFs alone; *cyan*, CAF cells in co-culture not making contacts with Panc1 cells; *blue*, CAFs making membrane contacts with Panc1 cells. Data are represented as mean ±SEM. ****, *p*<0.0001 as compared with CAFs making membrane contacts with Panc-1 cells (unpaired two-sided t-test). (**D**) Measurements of Ca^2+^ influx in CAFs using Fura2 fluorescent calcium indicator. CAFs were loaded Fura2-AM for 30mins and treated with vehicle, CM4620 5µM, or BTP2 1µM for 10mins. *Tg*, 2 μM thapsigargin was added to deplete ER Ca^2+^ stores in the absence of Ca2+. Once Fura2 signal returned to the baseline, Ca^2+^ was elevated to 1 mM. (**E**) Estimated SOCE in CAFs. The magnitude of SOCE was assessed by subtracting basal F340/F380 from the maximum F340/F380 in CAF cells treated with vehicle, CM4620, or BTP2. (**F**) Total annexin XII fluorescence intensity measured by microscopy in CAF-Panc-1 co-cultures in the presence of vehicle or Orai channel inhibitor BTP2 (10 µM), or clofazimine (1 µM). *Right*, representative images of annexin XII labeling. Data of individual images obtained in two independent experiments are represented as mean ±SD. *, *p*<0.05; **, *p*<0.01 (unpaired two-tailed t-test). (**G**) Polarization of MAPPER expression at the CAF-Panc-1 interface. Representative reconstituted (maximum intensity) confocal Z-stack image of DsRed-tagged Panc-1 and CAF cells transfected with GFP-MAPPER in co-culture. (**H**) Representative images of absence of polarization of ANO6 and MAPPER at the CAF-CAF contact sites. (**I**) Mean fluorescence intensity (MFI) of MAPPER and ANO6 comparing contact sites with cell body away from the contacts are shown as mean ±SD. ns, *p*>0.05 (unpaired two-tailed t-test). (**J**) Representative images of absence of polarization of ANO6 and STIM1 at the CAF-CAF contact sites. In E and F, CAFs were co-transfected either with ANO6-mCherry and GFP-MAPPER or ANO6-mCherry and YFP-STIM1. Fluorescence distribution of ANO6, Mapper, and STIM1 near the cell contact site (*magenta*) or away from the contact site (*white*) is shown by boxes. (**K**) Mean fluorescence intensity (MFI) of STIM1 and ANO6 comparing contact sites with cell body away from the contacts are shown as mean ±SD. ns, *p*>0.05 (unpaired two-tailed t-test).

**Supplementary Figure 5 related to Figure 5. ANO6 regulates cholesterol metabolism and tumor-promoting function of CAFs. (A)** Cell area of ANO6-KD fibroblasts relative to control CAFs (CAF1 line). Data represents mean ± SEM of cell area measured using Image J of phase contrast images from 5 in dependent cultures; **, *p*<0.01 (unpaired two-tailed t-test). (**B**) Quantification of Nile red fluorescence. Data are represented as mean±SD fluorescence intensity relative to controls obtained from 4 or more technical replicates in 3 independent experiments. ***, *p*<0.001; ****, *p*<0.0001 (unpaired two-sided t-test). (unpaired two-sided t-test). (**C**) Quantification of Nile red fluorescence of lipid droplets in control CAF1 cells, ANO6-KD, or ANO6-KD cells modified with a to lentivirus express the wild-type ANO6-mCherry, or ANO6 with indicated mutations (D409G, D703R, and Y563A). Data are represented as mean ±SD of 5 technical replicates combined from 2 independent experiments. **, *p*<0.01; ***, *p*<0.001; ****, *p*<0.0001 (unpaired two-sided t-test). (**D**) Viability of cholesterol auxotroph Panc-1^_NSDHL^ DsRed-tagged cancer cells in lipid-depleted serum (5% LDS) co-cultured with control CAF1 cells or ANO6-KD fibroblasts. Data are presented as mean ± SEM of ratio of dead (SYTOX Blue-positive) and live (DsRed-positive) Panc-1 cells. Fluorescent images were acquired on automated high throughput ImageXpress (Molecular Devices) microscope and analyzed using MetaXpress software. (**E**) PtdSer externalization induced by addition of ionomycin (10 µM, 5 minutes) or under basal culture conditions (**F**) assessed by intravital annexin XII staining of CAF membranes of control CAF1 cells, ANO6-KD, or ANO6-KD plus wild-type murine ANO6-mCherry (calcium-responsive), or with indicated modifications: D409G (constitutively active), D703R (inactive due to calcium-domain mutation), and Y563A (constitutively active). Graph represents total annexin XII fluorescence intensity represented as mean ±SD. ***, *p*<0.001, ****, *p* <0.0001 as compared with control CAFs treated with ionomycin (unpaired two-tailed t-test). (**G**) CRIPSRi-mediated silencing of NADP-sterol-dehydrogenase-like (NSDHL) in Panc-1 PDAC cells results in accelerated tumor growth. Representative images of pancreatic tumors at 5 weeks post implantation of 10^5^ of cancer cells to pancreatic tails of C.B-17.icr SCID mice. (**H**) Weights of CRISPRi-control and NSDHL-KD Panc-1 tumors as in (J). Data are presented as mean ± SD from n=6 tumors per condition; ***, *p*<0.001 (unpaired two-tailed t test).

**Supplementary Figure 6 related to Figure 6. Blockade of ANO6 restrains growth and blocks delivery of exogenous cholesterol to pancreatic tumors.** (**A, B**) Images of individual pancreatic tumors collected at 3 weeks post implantation of 10^5^ of KPC3 PDAC cells to pancreatic tails. Animals were treated with niclosamide (**A**) or clofazimine (**B**) as indicated in Fig. 6A.

**Supplementary Figure 7 related to Figure 7. Assessment of CAF-expressed ANO6 on the status of the cytotoxic T lymphocytes. (A)** Gating strategy for assessment of intratumoral and splenic immune cells (equally applied to all samples). Single cells in dissociated suspensions of pancreatic tumors were identified using combined SSC and FSC filtering to exclude debris, apoptotic cells and cell doublets as outlined with gates P1-P3. Both CD45-and CD45+ cells were present in the single cell populations. The analyses of immune cells were done exclusively on CD45+ population. “Backgating” was used to confirm the identified immune cell subtypes were associated with non-apoptotic single-cell populations. (**B**) Clofazimine has no effect on preponderance of immune subsets in the spleens of tumor-bearing mice. (**C**) Clofazimine does not change the percentage of intratumoral CD11b+F4/80+ macrophages, LY6G+ granulocytes and LY6C+ MDSCs. (**D**) Flow cytometry profiles of Filipin III staining corresponding to cholesterol levels in the splenic CD3+CD8+ CTLs from mice treated with clofazimine or vehicle as described in Fig. 6C. (**E**) Clofazimine has no effect on expression of activity (CD69) and exhaustion (PD-1 and TIM3) markers, or intracellular interferon-γ in splenic CD3+CD8+ CTLs isolated from mice treated with clofazimine or vehicle as described in Fig. 6C. (**F**) Flow cytometry profiles of activity (CD69) and exhaustion (PD-1 and TIM3) markers on the intratumoral CTLs from mice treated with clofazimine or vehicle as described in Fig. 6C.

**Supplementary Figure 8 related to Figure 7. Assessment of CAF-expressed ANO6 on the status of the cytotoxic T lymphocytes in vitro. (A)** Flow cytometry dot plots of DiD uptake by OT-1 CTLs cultured for 12 hours either alone or with DiD-labeled CAFs at 1:5 ratio in presence of niclosamide, clofazimine, annexin V, anti-TIM3 and 25HC, respectively. (**B**) Flow cytometry profiles of Filipin III of OT-1 CTLs cultured for 12 hours either alone or with DiD-labeled CAFs at 1:5 ratio in presence of niclosamide, clofazimine, annexin V, anti-TIM3 and 25HC, respectively. (**C**) Flow cytometry dot plots of intracellular IFN-γ in OT-1 CTLs co-cultured for 12 hours either alone or with DiD-labeled CAFs at 1:5 ratio in presence of niclosamide, clofazimine, annexin V, anti-TIM3 and 25HC, respectively. (**D**) Flow cytometry dot plots of DiD uptake by OT-1 CTLs cultured for 12 hours either alone or with DiD-labeled control CAFs or ANO6-KD CAFs. (**E**) Flow cytometry profiles of Filipin III in OT-1 CTLs cultured for 12 hours either alone or with control CAFs or ANO6-KD CAFs. (**F**) Flow cytometry contour density plots of intracellular IFN-γ in OT-1 CTLs cultured for 12 hours either alone or with control CAFs or ANO6-KD CAFs. In all graphs, data are presented as mean ± SEM from n=5 tumors per treatment or n=4 replicates for each in vitro condition; *p*-values were calculated by unpaired two-tailed t test: *, *p*<0.05; **, *p*<0.01; ***, *p*<0.001; ****, *p*<0.0001.

## Supplementary videos

<u>Supplementary video 1</u>. **Intravital imaging of BODIPY-cholesterol-labeled LDL particles immediately following intravenous injection**. BODIPY-cholesterol fluorescence (*green*) carried in 25 nm-sized LDL particle can be seen in the peritumoral blood vessels immediately after i.v. injection. Note the absence of green fluorescence in DsRed-positive PDAC tumor cells. Images were obtained on Leica Microsystems multiphoton microscope with 25x objective.

<u>Supplementary video 2.</u> **Intravital imaging of orthotopic tumors at xx-XX hours following intravenous injection of DiI-labeled LDL particles.** Migration of GFP-positive KPCN349 PDAC tumor cells (*green*) engaging via direct contacts the stromal cells saturated with LDL (*orange*). Images were obtained on Leica Microsystems multiphoton microscope with 25x objective.

<u>Supplementary video 3.</u> **Enhanced 3D reconstruction of confocal Z-stack images of CAF (green) and KPCN349 (pink) interface taken at 2-second intervals.** At the site of membrane contacts, extensive CAF plasma membrane blebbing (*green*, BODIPY-cholesterol) projecting to the cytoplasm of carcinoma cells (*pink*) is observed. Confocal laser microscopy with 60x objective. *Scale bar*, 2 µm.

<u>Supplementary video 4</u>**. Phosphatidylserine externalization on CAF membrane is induced by cell-cell contacts with cancer cells.** CAFs co-cultured with DsRed-tagged Panc-1 cells in the presence of Annexin XII (*green*) and propidium iodide (pSIVA, Abcam). Confocal laser microscopy with 60x objective.

<u>Supplementary video 5</u>**. Lack of phosphatidylserine externalization in ANO6-KD CAFs in co-culture with cancer cells.** CAFs co-cultured with DsRed-tagged Panc-1 cells in the presence of Annexin XII (*green*) and propidium iodide (pSIVA, Abcam). Confocal laser microscopy with 60x objective.

<u>Supplementary video 6.</u> **Fluorescence of cytosolic Ca^2+^ indicator Fluo4 in cocultures of DsRed-tagged Panc-1 and CAFs.** CAF and Panc-1 cells in co-culture on coverslips were pre-loaded with the calcium indicator (*green*, 5 µM Fluo4-AM), washed, incubated for additional 30 minutes and imaged for 60 cycles at a frequency of 0.33 Hz using a Leica SP8 laser scanning microscope (Buffalo Grove, IL) with 60x objective.

<u>Supplementary video 7</u>. **Blocking Orai channels with BTP2 abrogates calcium entry induced via CAF-Panc-1 cell contacts**. The settings are as in the Supplementary video 6.

<u>Supplementary video 8.</u> **Calcium-induced membrane ruffling in CAFs is ANO6-dependent.** CRISPRi-control CAFs were imaged after addition of 10 µM ionomycin for 5 minutes and imaged using intravital label-free interferometry-based microscopy (Nanolive, 60x objective).

<u>Supplementary video 9</u>. **Absence of calcium-induced membrane ruffling in ANO6-deficient CAFs.** The settings are as in the Supplementary video 8.

## Acknowledgements.

We dedicate this work to Patricia Keely (pioneer in the field of ECM biology) and Neelima Shah (the most beautiful soul) who continue to inspire our research. We thank Dr. Celeste Simone and her team at the University of Pennsylvania for insightful discussions throughout this study. This project was supported by NIH core grant No. CA06927 in support of the following Fox Chase Cancer Center’s Facilities: High throughput, Animal, Histopathology, Cell Culture, Biosample Repository, Imaging, and Biostatistics. Support was also provided by the Pew Charitable Fund, the 5^th^ AHEPA Cancer Research Foundation, and by a generous gift from Mrs. Concetta Greenberg to the M&C Greenberg Pancreatic Cancer Institute at Fox Chase Cancer Center. Authors were also supported by NIH S10ODO23666; by R01 CA188430, and R21 CA164205, (to I. Astsaturov); R21 CA231252 (to I. Astsaturov and E. Cukierman; R01 CA232256, and R01 CA269660 (to E. Cukierman); K99 GM148819 (to JC. Gardiner and mentor E. Cukierman); R01 CA240814 (to S.Y.Fuchs). Further funds used were from the DOD W81XWH-22-PCARP-IDA (to DB. Vendramini-Costa and E. Cukierman); the Emerald Foundation’s Black in Cancer Award and ACS’s PF-18-218-01-CSM (to JC. Gardiner); two Pancreatic Cancer Action Network (PanCan) grants, the Career Development Award in memory of Skip Viragh (to R. Francescone), and the Postdoctoral Fellowship intended for a French National trainee (to C. Ogier); the French Labellisation program of “Ligue Contre le Cancer” and of “Institut National du Cancer INCa_18389” (to C. Bousquet), a grant from The William Wikoff Smith Foundation (to E.A.Golemis).

