## Supplementary figures and images for "Trogocytosis of cancer-associated fibroblasts promotes pancreatic cancer growth and immune suppression via phospholipid scramblase anoctamin 6 (ANO6)"

### Supplemental Figure 1

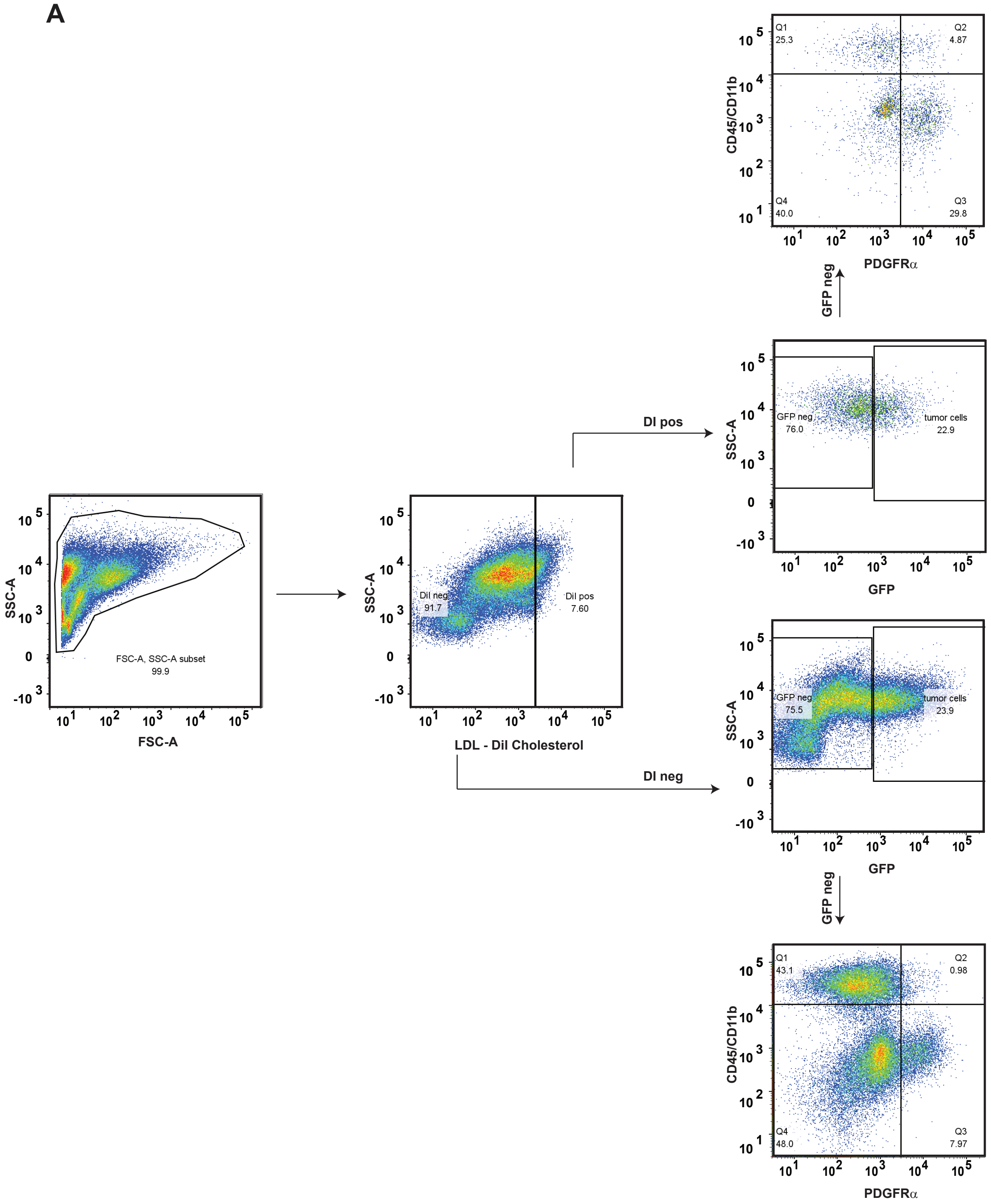

### Supplemental Figure 2

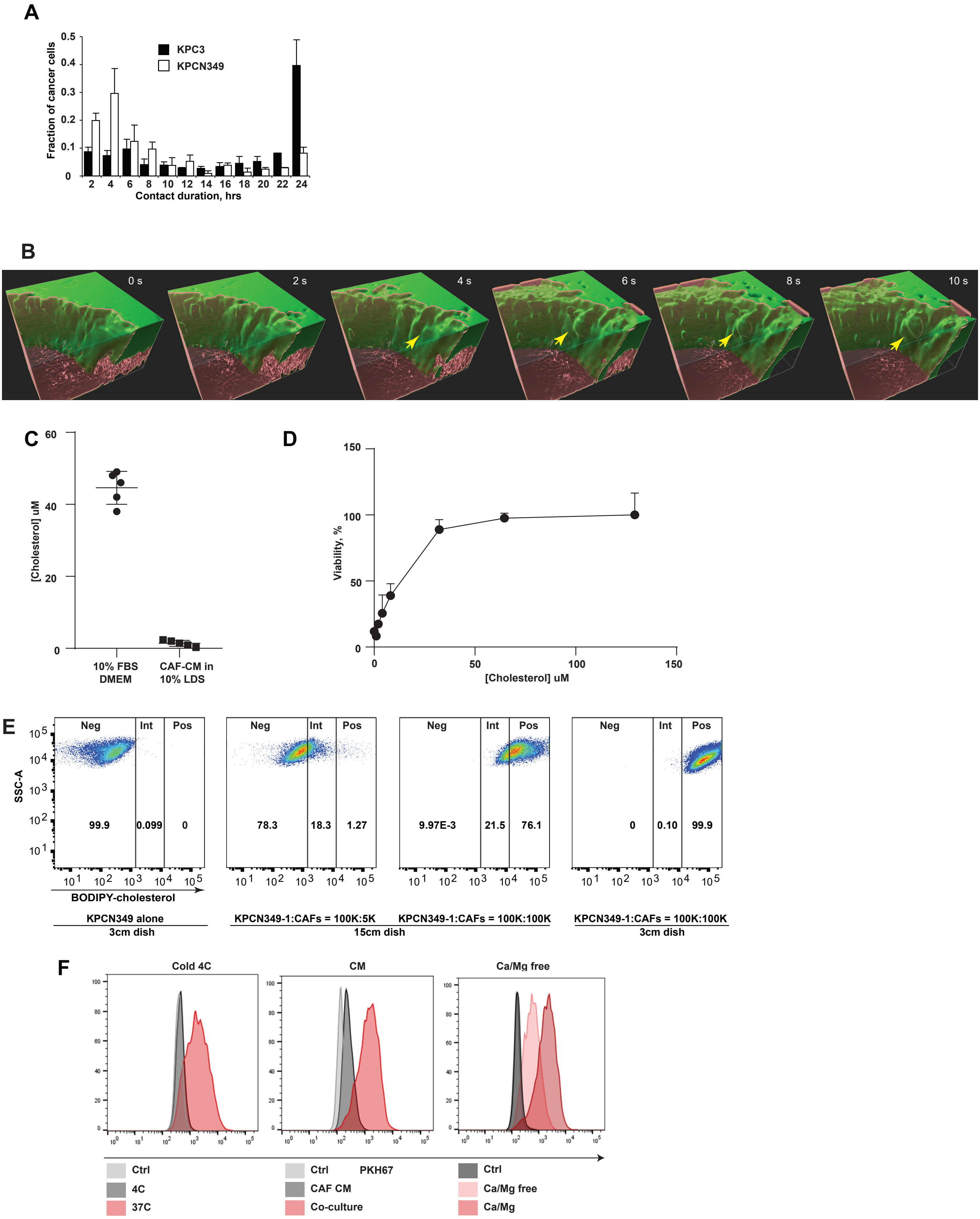

### Supplemental Figure 3

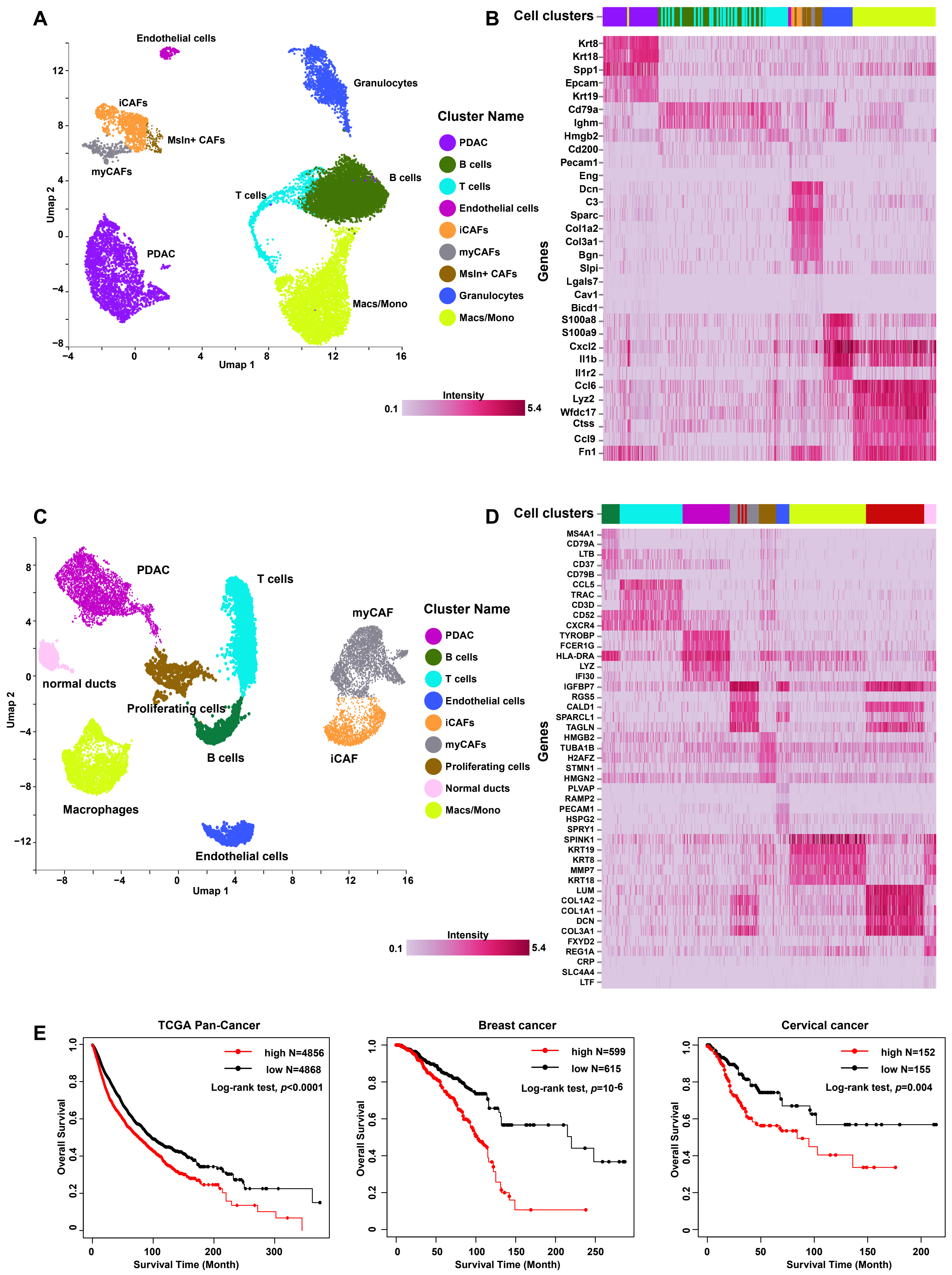

### Supplemental Figure 4

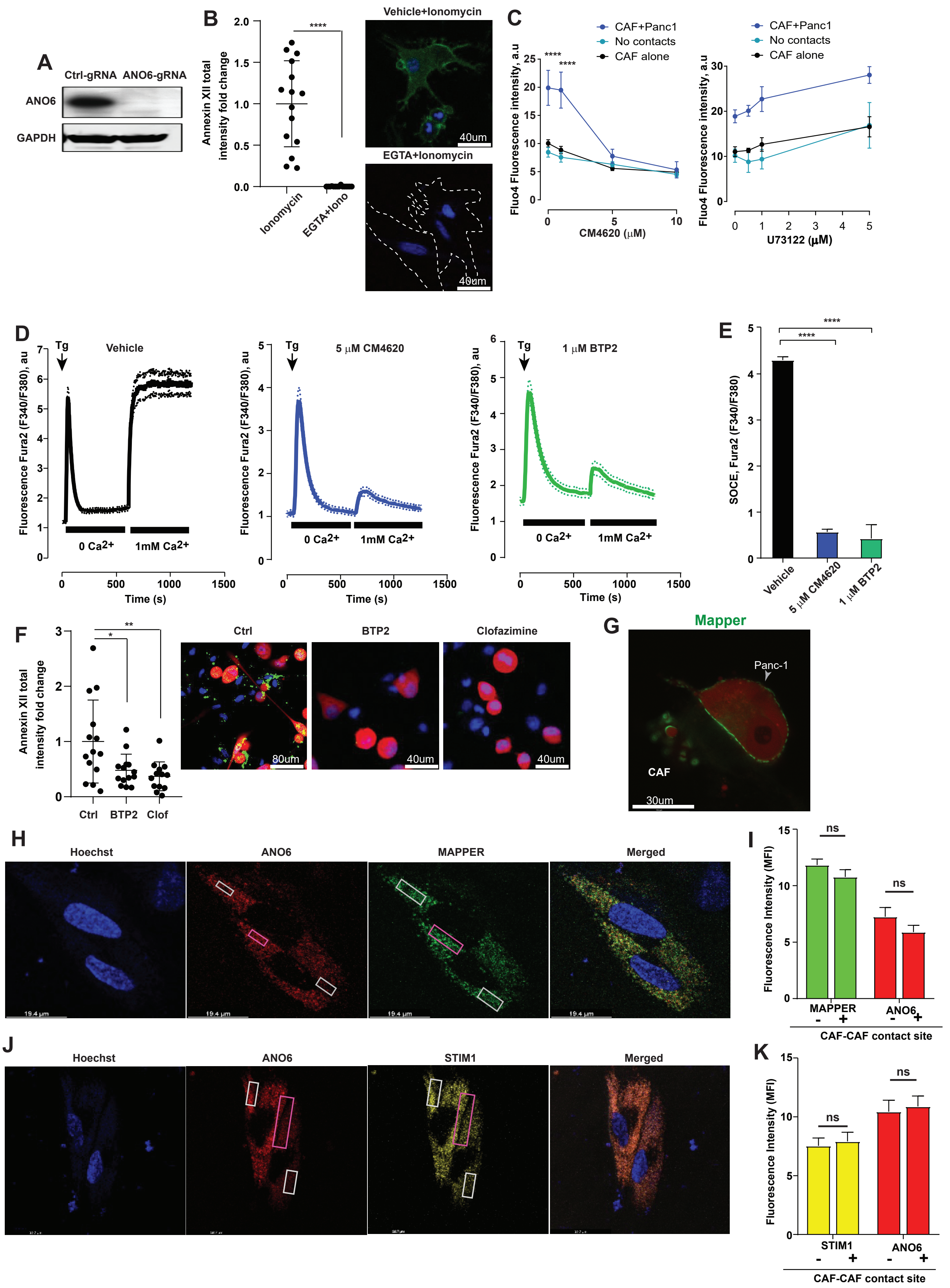

### Supplemental Figure 5

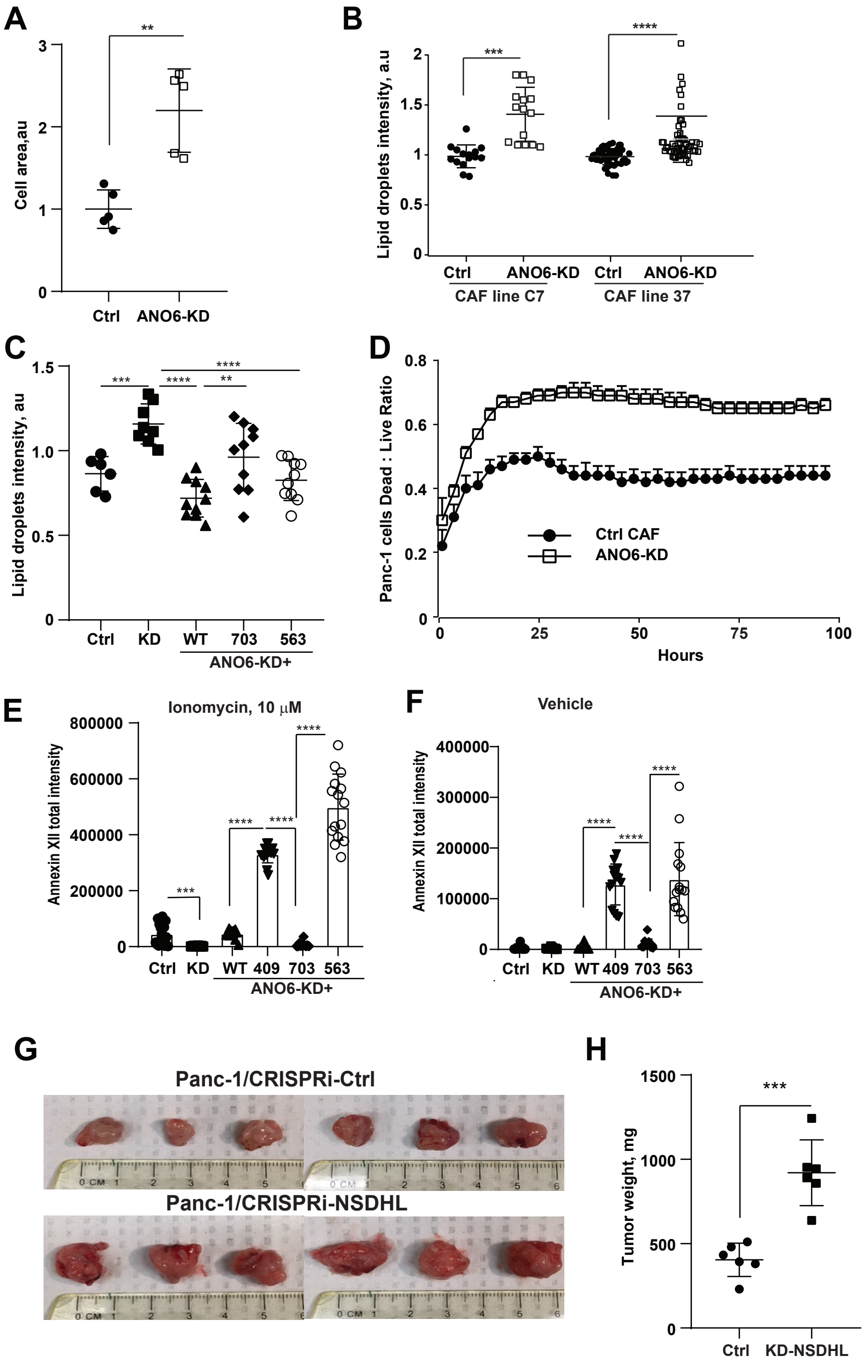

### Supplemental Figure 6

**Supplementary Figure 6**

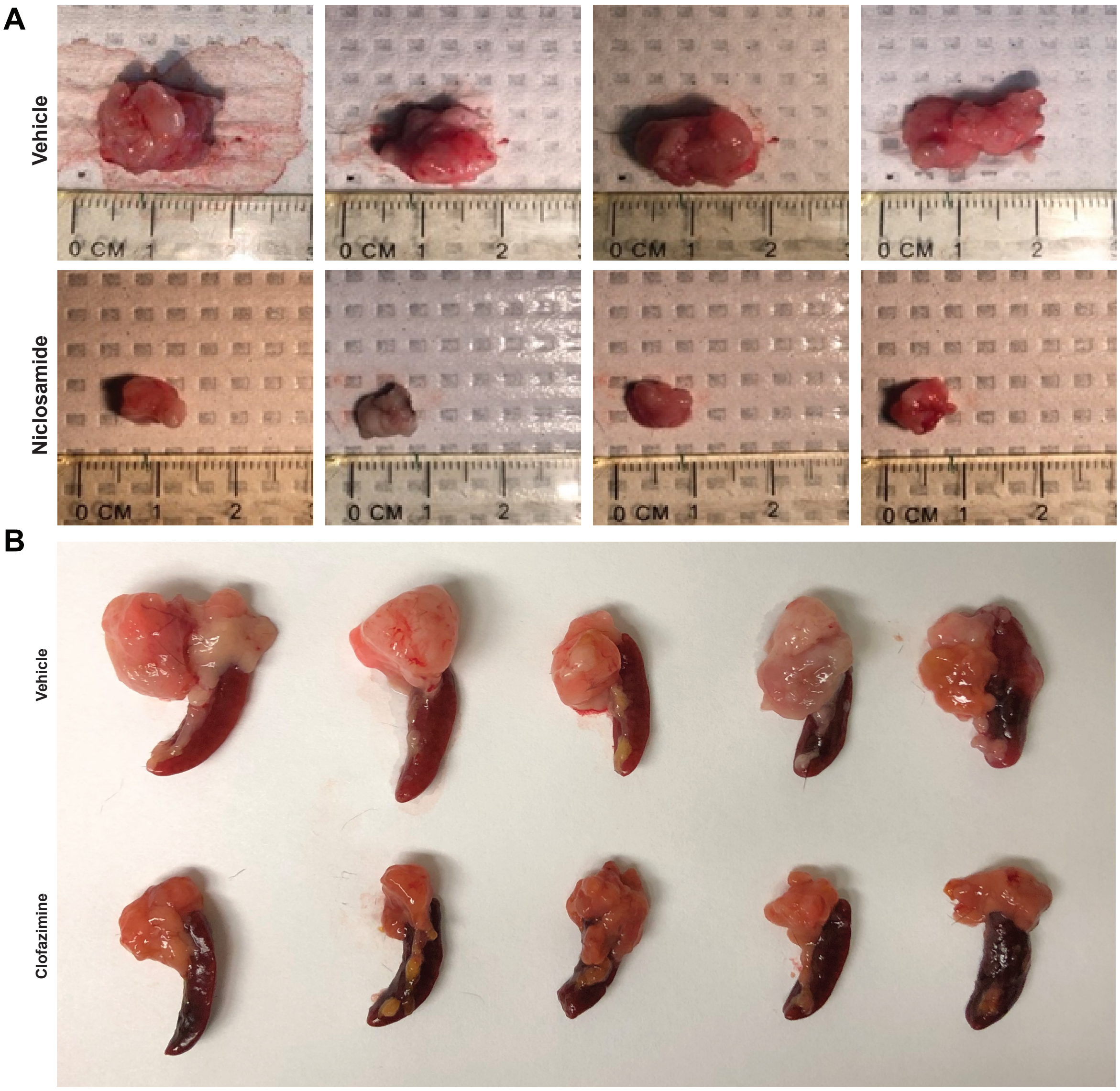

### Supplemental Figure 7 and 8

#### Supplementary Figure 7

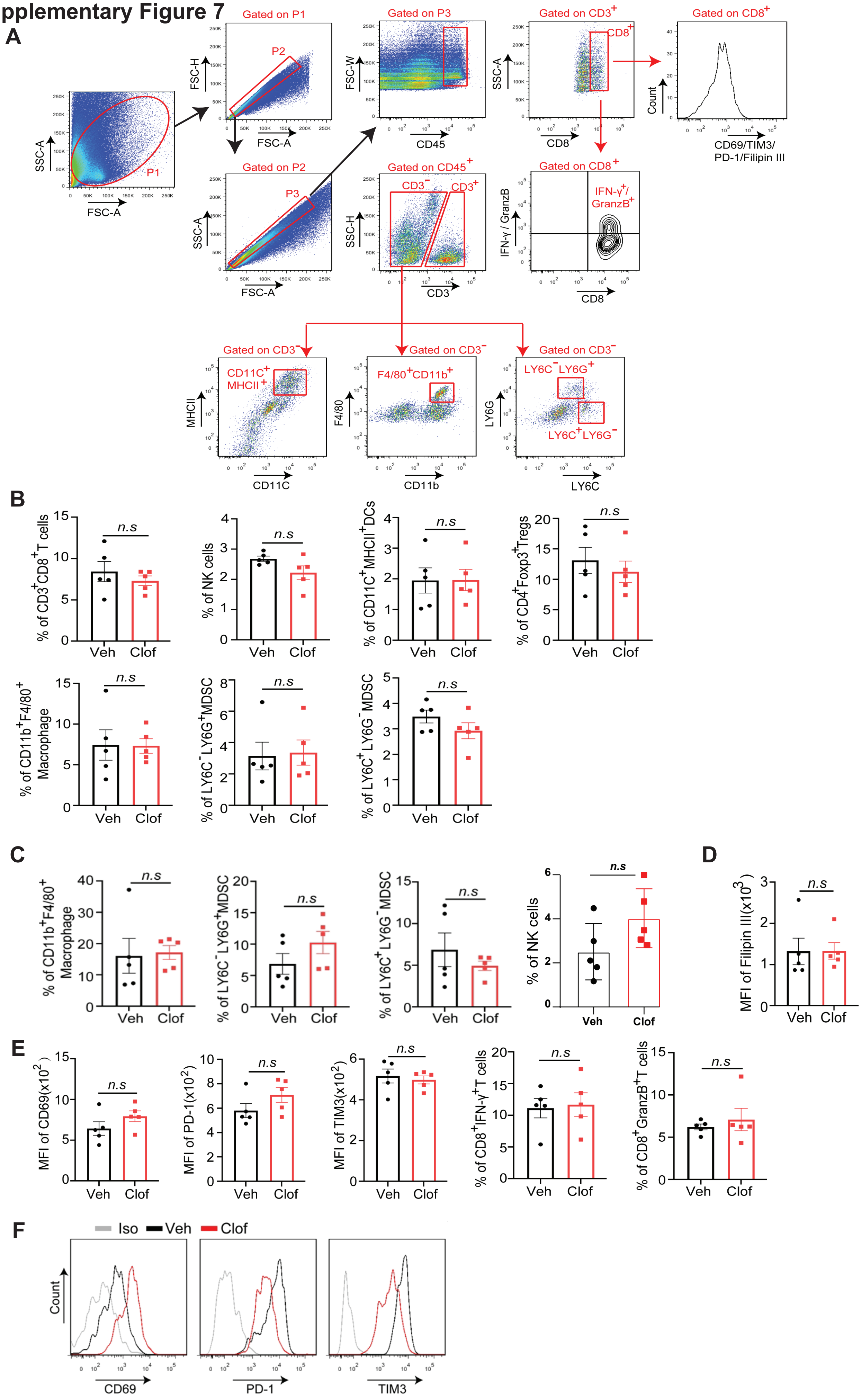

#### Supplementary Figure 8

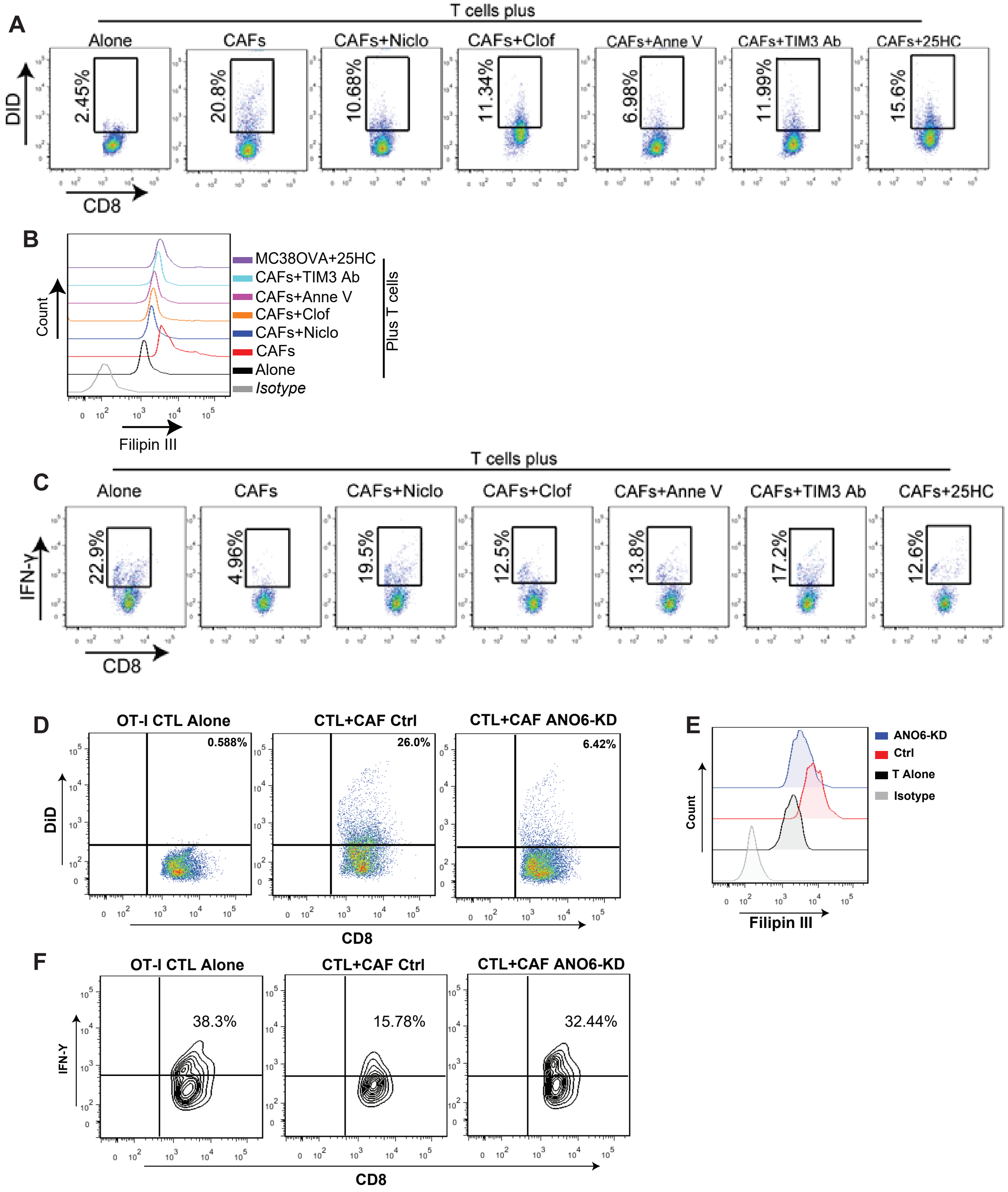
